# Actin binding domain of Rng2 sparsely bound on F-actin strongly inhibits actin movement on myosin II

**DOI:** 10.1101/2020.04.14.041046

**Authors:** Yuuki Hayakawa, Masak Takaine, Kien Xuan Ngo, Taiga Imai, Masafumi D. Yamada, Arash Badami Behjat, Kenichi Umeda, Keiko Hirose, Ayhan Yurtsever, Noriyuki Kodera, Kiyotaka Tokuraku, Osamu Numata, Takeshi Fukuma, Toshio Ando, Kentaro Nakano, Taro Q.P. Uyeda

## Abstract

Substoichiometric binding of certain actin-binding proteins induces conformational changes in a disproportionally large number of actin protomers in actin filaments. Here, we report a case in which such conformational changes in actin filaments have profound functional consequences. Rng2 is an IQGAP protein implicated in the assembly and contraction of contractile rings in *Schizosaccharomyces pombe*. We found that the calponin-homology actin-binding domain of Rng2 (Rng2CHD) strongly inhibits the motility of actin filaments on myosin II *in vitro.* On skeletal muscle myosin II-coated surfaces, Rng2CHD halved the sliding speed of actin filaments at a binding ratio of 1.3% (=1/77), and virtually stopped movement at a binding ratio of 11% (=1/9). Rng2CHD also inhibited actin movements on *Dictyostelium* myosin II, but in this case by inducing the detachment of actin filaments from myosin II-coated surfaces. Rng2CHD induced cooperative structural changes of actin filaments accompanied by shortening of the filament helical pitch, and reduced the affinity between actin filaments and subfragment 1 (S1) of muscle myosin II in the presence of ADP. Intriguingly, actin-activated ATPase of S1 was hardly inhibited by Rng2CHD. We suggest that sparsely bound Rng2CHD induces global structural changes of actin filaments and interferes with the force generation by actin-myosin II.

## Introduction

Actin exists in all eukaryotic cells and performs a wide variety of functions. The interaction between actin and myosin II drives a variety of motile activities such as muscle contraction (Huxley and Niedergerke, 1954; Huxley and Hanson, 1954), the amoeboid movement of cells (Pollard et al., 1974; Clarke and Spudich, 1977; Korn, 1978), and the contraction of contractile rings (CRs). A CR is a ring-like structure that appears transiently on the equatorial plane underneath the cell membrane during cytokinesis of animal cells and many unicellular eukaryotes (Pollard, 2010). A CR consists of actin filaments and myosin II filaments, together with a number of actin-binding proteins (ABPs). It is thought that a CR contracts by sliding between the two filament systems due to actomyosin movement (Mabuchi and Okuno, 1977; Satterwhite and Pollard, 1992). The mechanism of assembly/disassembly of CRs, as well as the mechanism by which contraction is regulated, have been subjected to extensive research, and recent advances using various model organisms, particularly the fission yeast *Schizosaccharomyces pombe*, have established the roles of major factors in these processes (Goyal et al., 2011). However, CRs are under elaborate spatiotemporal control involving a plethora of ABPs and other regulatory proteins, and our understanding of their functions is far from complete.

One such yet incompletely characterized CR-related ABP is Rng2 in *S. pombe*. Rng2 is an IQ motif-containing GTPase activating protein (IQGAP) that plays important roles in the formation of CRs (Eng et al., 1998; Takaine et al., 2009). It is believed that Rng2 crosslinks and bundles actin filaments together to form CRs because only abnormal accumulation of actin filaments was observed at their division sites in Rng2 knockout *S. pombe* cells and in temperature-sensitive Rng2 mutant cells at a restrictive temperature (Eng et al., 1998; Takaine et al., 2009). Additionally, temperature-sensitive Rng2 mutant cells showed normal actin filament rings at the permissive temperature but the distribution of myosin II on actin filaments was abnormal (Takaine et al., 2009). This latter result suggests that Rng2 regulates the interaction between actin filaments and myosin II filaments. Therefore, we have been investigating if Rng2CHD, a 181 amino acid residue-long calponin-homology actin binding domain (CHD) at the N-terminus of Rng2, exhibits *in vitro* activities that are related to the regulation of assembly and/or contraction of CRs.

To our surprise, we found that Rng2CHD inhibits, rather than promotes, actomyosin motility during *in vitro* motility assays in which actin filaments move on surfaces coated with skeletal muscle myosin II or *Dictyostelium* non-muscle myosin II. We investigated the mechanism of inhibition using muscle myosin II, which was more strongly inhibited by Rng2CHD. We found that Rng2CHD bound on actin filaments strongly inhibited actomyosin II motility even when binding was sparse, without significantly inhibiting actin-activated myosin ATPase activity. Sparsely bound Rng2CHD induced cooperative conformational changes in actin filaments, and those actin filaments displayed a reduced affinity for the motor domain of myosin II carrying ADP, demonstrating that this is a novel mode of regulation of actomyosin II motility.

## Results

### Rng2CHD strongly inhibits sliding of actin filaments on myosin II *in vitro*

To examine if Rng2 stimulates or inhibits actomyosin motility, we performed *in vitro* motility assays in which actin filaments move on a glass surface coated with myosin II (Kron and Spudich, 1986). We examined two types of myosin II, skeletal muscle myosin II and non-muscle myosin II derived from *Dictyostelium discoideum*. The movement of actin filaments was significantly inhibited by Rng2CHD on both types of myosin II (Figure 1). However, the apparent mode of inhibition was different between the two systems. On surfaces coated with full-length *Dictyostelium* myosin II, sliding velocity in the presence of 1 mM ATP was not significantly slowed by 100-500 nM Rng2CHD, but a large fraction of actin filaments dissociated from the myosin-coated surface in the presence of 1 µM Rng2CHD (Figure 1B and Video 1). Trajectory analyses also showed that actin filaments moving on *Dictyostelium* myosin II in the presence of Rng2CHD tended to slide sideways (Figure 1C), a phenomenon typically observed in standard *in vitro* motility assays when the surface density of myosin motor is too low.

**Figure 1.**
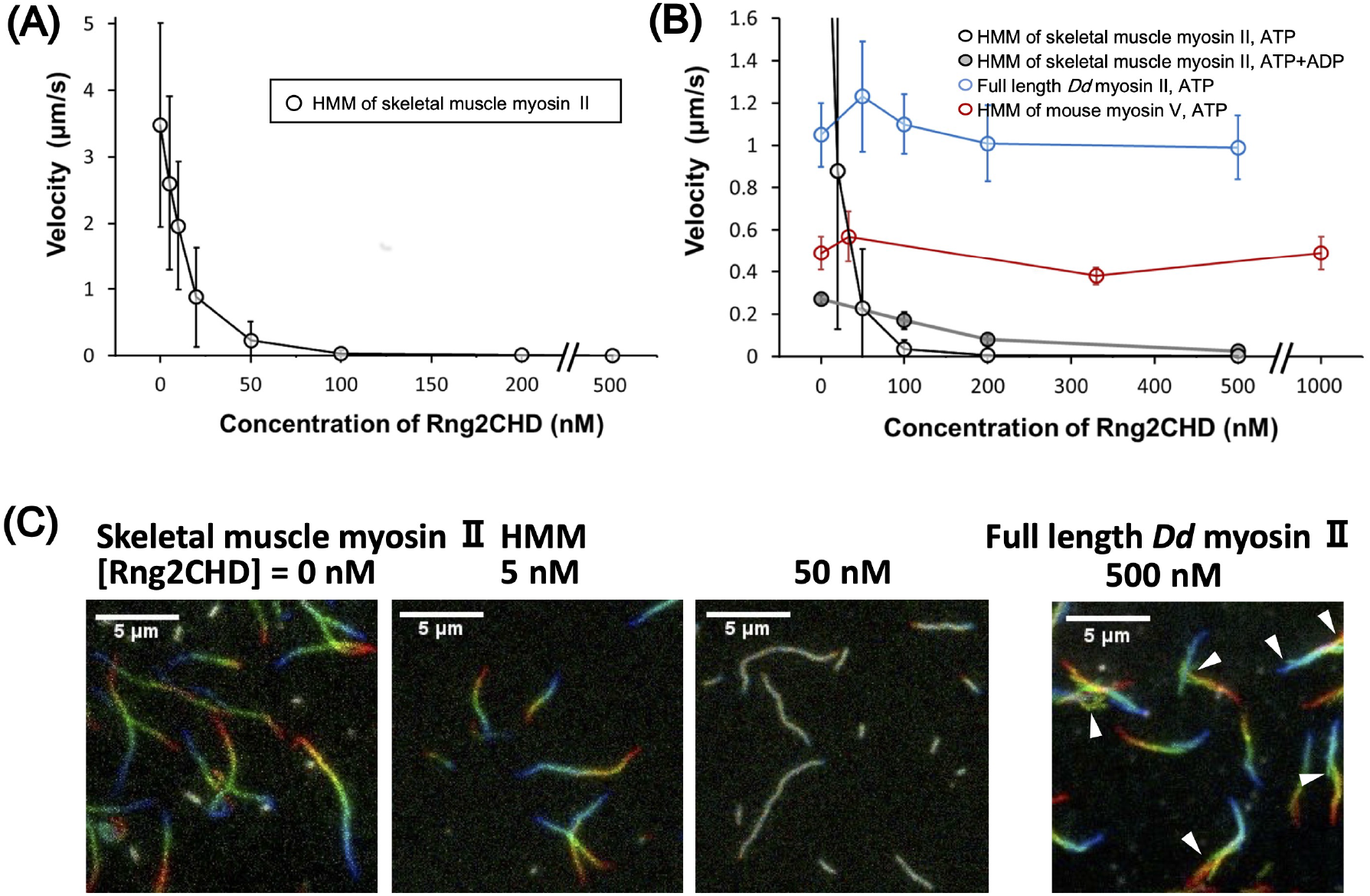
Rng2CHD strongly inhibits movement of actin filaments on myosin II, but not on myosin V. **(A)** Movement velocity of actin filaments on muscle myosin II HMM in the presence of various concentrations of Rng2CHD. **(B)** Velocity of actin filaments on surfaces coated with muscle HMM, full length *Dictyostelium* myosin II and myosin V HMM in the presence of various concentrations of Rng2CHD. Open black circles show speeds of actin filaments by muscle HMM in the presence of 1 mM ATP, and black circles filled with gray in the presence of 0.2 mM ATP and 1 mM ADP. Open blue circles show speeds by *Dictyostelium* myosin II in the presence of 1 mM ATP, and red open circles show speeds by myosin V in the presence of 2 mM ATP. A total of randomly chosen 100-110 filaments, excluding very short filaments (<1.5 µm), were analyzed for each condition. Data are expressed as the mean ± SD. For other methods of velocity analyses, see Figure Supplement 1. The movement velocity on muscle HMM in the presence 100 nM Rng2 CHD was 0.035 ± 0.043 µm/s in the presence of 1 mM ATP and 0.17 ± 0.04 µm/s in the presence of 0.2 mM ATP and 1 mM ADP, and this difference was statistically significant (*p*<10^-37^ by Student’ *t*-test). In the presence of 200 nM Rng2CHD, the two velocities were 0.0057 ± 0.0043 µm/s and 0.079 ± 0.024 µm/s, respectively (*p*<10^-21^). Actin velocity on *Dictyostelium* myosin II-coated surfaces was 1.1 ± 0.2 µm/s in the absence of Rng2CHD and 1.2 ± 0.3 µm/s in the presence of 50 nM Rng2CHD. This difference was statistically significant (*p*<10^-13^ by Student’s *t*-test) and was reproduced in two independent experiments. For a possible explanation for this small difference, see Supplementary information 2. In the presence of 1 µM Rng2CHD, most of the actin filaments detached from the surfaces coated with *Dictyostelium* myosin II, after brief unidirectional movements (Video 1). **(C)** Trajectories of moving actin filaments on muscle myosin II HMM in the presence of 0, 5 and 50 nM Rng2CHD, and on *Dictyostelium* myosin II in the presence of 500 nM Rng2CHD. Eleven consecutive images at 0.2 s (muscle HMM) or 0.5 s (*Dictyostelium* myosin II) intervals are coded in rainbow colors from red to blue, and overlaid. Note that actin filaments moving on *Dictyostelium* myosin II in the presence of Rng2CHD often move laterally, indicating weak affinity between actin filaments and the myosin motors (white arrowheads). Because filament ends also frequently move laterally, mean actin velocities on *Dictyostelium* myosin II in the presence of Rng2CHD, which were calculated by tracking filament ends, were overestimated.

On surfaces coated with heavy meromyosin (HMM) of rabbit skeletal muscle myosin II, in contrast, actomyosin motility in the presence of 1 mM ATP was significantly slowed in the presence of low concentrations of Rng2CHD. Movements were completely stalled, and all the filaments were virtually immobilized to the surface in the presence of 200 nM Rng2CHD (Figure 1A and Video 2). Trajectory analyses showed that under all the inhibition conditions tested using muscle HMM, the front end of the actin filament was followed by the remainder of the filament (Figure 1C). Buckling of the filaments, indicative of local inhibition of movement, was rarely observed. This indicates that movement is more or less uniformly inhibited along the entire length of the filaments on muscle HMM surfaces. Movements of actin filaments on surfaces deposited with filaments of muscle myosin II were similarly inhibited by Rng2CHD, indicating that the inhibition is not related to the mode of immobilization of myosin motors (Video 3).

The two myosin IIs differ not only in their response to Rng2CHD, but also in their sliding velocity in the absence of Rng2CHD. The latter difference can be primarily attributed to a difference in the lifetime of the force-generating A•M•ADP complex (Toyoshima et al., 1990; Uyeda et al., 1990). Indeed, previous kinetic measurements demonstrated that the dissociation of *Dictyostelium* myosin II motor carrying ADP from actin is about 5-fold slower and ATP-induced dissociation of actin-myosin motor complexes is 10-fold slower than muscle myosin II (Ritchie et al., 1993). Thus, we examined the effects of extending the lifetime of the force-generating complex of actin and muscle HMM carrying ADP (Figure 1B). In the presence of 0.2 mM ATP and 1 mM ADP, the sliding velocity by muscle HMM in the absence of Rng2CHD was slowed to 0.27 µm/s, which was expected, but was further slowed only by an additional 37% by 100 nM Rng2CHD, in contrast to 99% inhibition by the same concentration of Rng2CHD in the presence of 1 mM ATP. Similar results were obtained in the presence of 200 nM Rng2CHD. Consequently, in the presence of 100 nM or 200 nM Rng2CHD, filaments moved faster in the presence of 0.2 mM ATP and 1 mM ADP than in the presence of 1 mM ATP (Video 4). Thus, extension of the A•M•ADP complex of muscle HMM partially mimicked the property of *Dictyostelium* myosin II in terms of sensitivity to Rng2CHD.

We also performed *in vitro* motility assays in which actin filaments moved on recombinant myosin V HMM that was expressed in insect cells. In striking contrast to myosin II, up to 1 µM Rng2CHD did not inhibit the sliding of actin filaments on myosin V HMM (Figure 1B and Video 5).

### Rng2CHD on actin filaments inhibits actomyosin II movement on muscle HMM *in vitro* even when binding is sparse

To characterize the inhibition of motility by Rng2CHD, we decided to use fragments of muscle myosin II in the following experiments since the movement by muscle myosin II was most strongly inhibited by Rng2CHD. First, we estimated the binding ratio, or molar binding density, of Rng2CHD to actin protomers when the movement of actin filaments was potently inhibited on muscle HMM. The concentrations of Rng2CHD that caused 50%, 75% and 95% reduction of movement speed, as estimated from the velocity curve, were 12 nM, 21 nM and 64 nM, respectively (Figure 1A). In parallel, we performed co-sedimentation assays of actin filaments with Rng2CHD, and the dissociation constant (*K_d_*) between Rng2CHD and actin protomers was calculated (Figure 2A, 2B). *K_d_* was determined to be 0.92 μM by the following fitting function:

**Figure 2.**
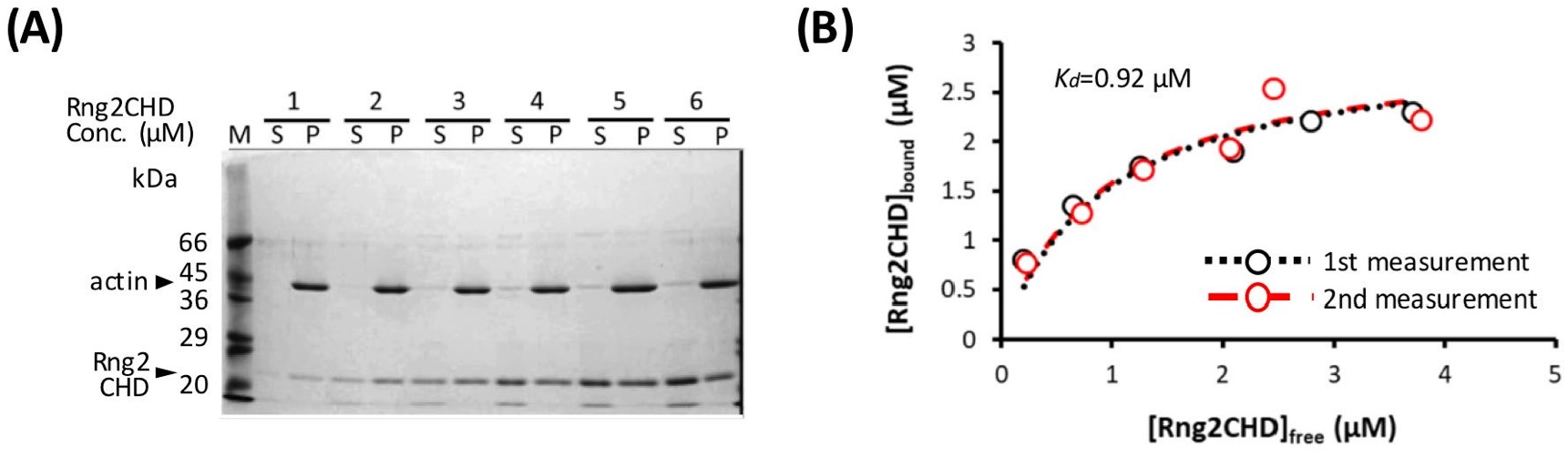
Measurement of *K_d_* between actin filaments and Rng2CHD. **(A)** Co-sedimentation assay of Rng2CHD with 3 µM of actin filaments. **(B)** The concentration of bound Rng2CHD was plotted against the concentration of the free fraction and fitted with the following equation: [*Rng2CHD_bound_*] = [*Actin_total_*] [*Rng2CHD_free_*] / ([*Rng2CHD_free_*+*K_d_*]). *K_d_* between Rng2CHD and actin protomers was calculated as an average value of the two trials.

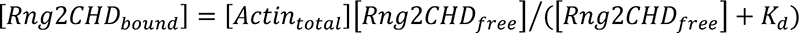 (Eq. 1 in Materials and Methods).

During *in vitro* motility assays, in which the concentration of actin protomers is much lower than that of Rng2CHD, it is difficult to estimate [*Actin_total_*], but the following approximation holds: 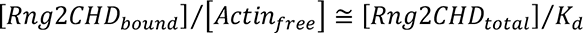 (Eq. 3 in Materials and Methods). Using this approximation, the binding ratio of Rng2CHD to actin protomers that caused 50%, 75% and 95% reduction of actomyosin II movement speed on muscle HMM was estimated to be 1.3%, 2.2% and 6.7%, respectively (Table 1). In other words, Rng2CHD was sparsely bound to actin filaments when the speed was reduced to half, and at the Rng2CHD concentration of 100 nM, when the mean sliding velocity was reduced to 1% of the control, the binding ratio was 11%.

**Table 1.** Estimated binding ratio of Rng2CHD and GFP-Rng2CHD to actin filaments.

| Inhibition rate of movement | 50% | 75% | 80% | 90% | 95% |
| --- | --- | --- | --- | --- | --- |
| $[Rng2CHD_{bound}]/[Actin_{total}]$ | 1.3% | 2.2% | 3.0% | 4.5% | 6.7% |
| $[GFP - Rng2CHD_{bound}]/[Actin_{total}]$ | nd | nd | 9.7% | 21% | nd |

The estimated binding ratio of Rng2CHD and GFP-Rng2CHD to actin protomers that caused various degrees of movement inhibition on muscle myosin II HMM. These values were estimated based on the following approximation: 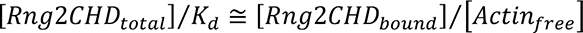 for Rng2CHD, and from fluorescence intensities for GFP-Rng2CHD (Figure 3). nd: not determined.

**Figure 3.**
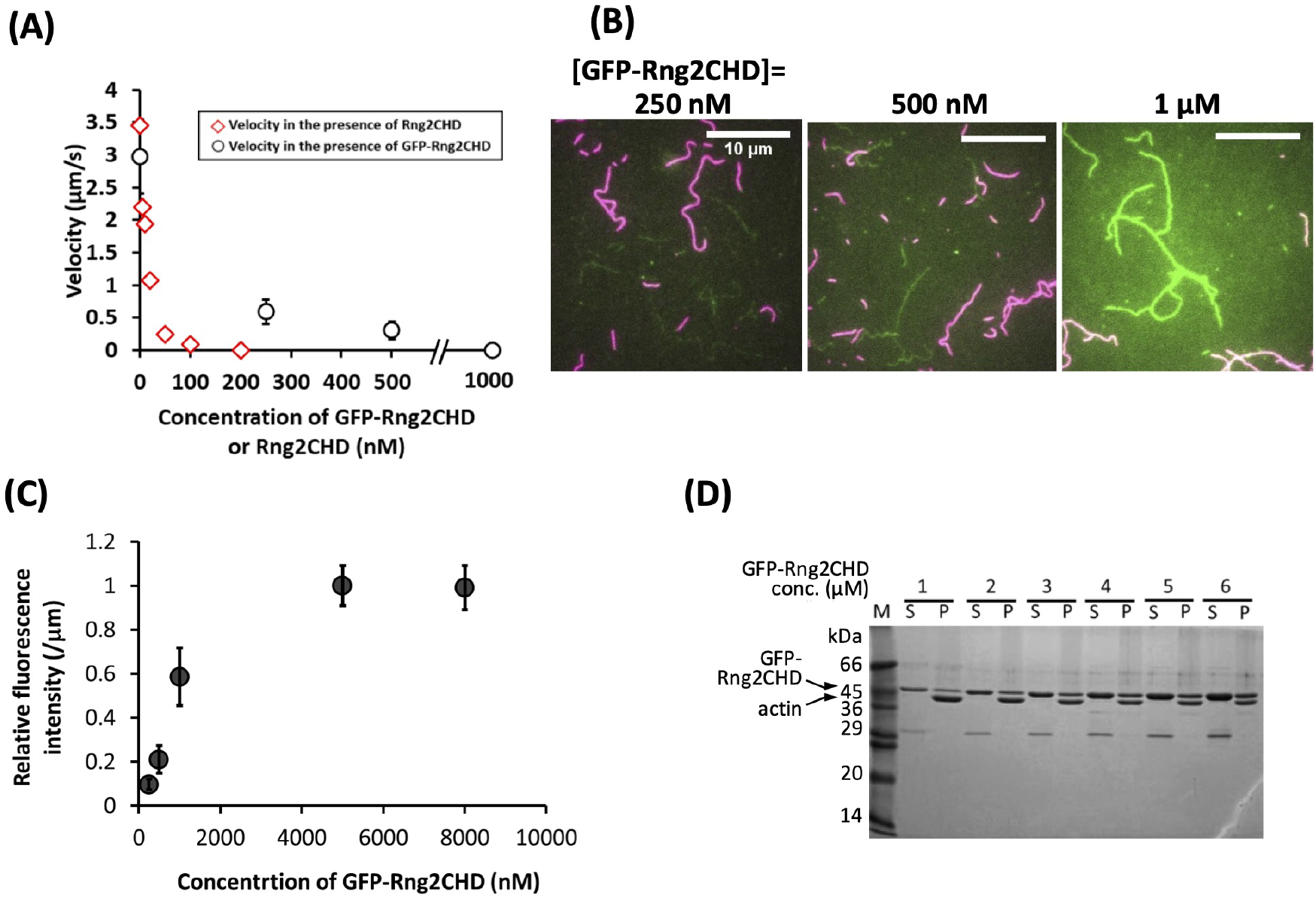
Sparsely bound GFP-Rng2CHD on actin filaments inhibits their movement on muscle myosin II HMM. **(A)** Movement velocity of actin filaments on muscle myosin II HMM-coated glass surfaces in the presence of various concentration of GFP-Rng2CHD (black plots). Ten smoothly moving filaments were chosen for each condition, and approximately 10 consecutive measurements were made for each filament. Data are expressed as the mean ± SD. Movement velocity in the presence of Rng2CHD is also shown for the reference (red plots). **(B)** Fluorescence micrographs of GFP-Rng2CHD bound to actin filaments. Actin filaments stabilized by non-fluorescent phalloidin and those labeled with rhodamine phalloidin were present at a 1:1 molar ratio. Green: GFP fluorescence. Red: rhodamine fluorescence. Non-fluorescent phalloidin-stabilized actin filaments were used to quantify the intensity of GFP fluorescence, since there was a low level of leakage of rhodamine fluorescence in the GFP channel, which could disturb the measurement of GFP fluorescence. Rhodamine-phalloidin–labeled actin filaments were used to measure sliding speed. Bars: 10 µm. **(C)** Fluorescence intensity of GFP-Rng2CHD on non-fluorescent phalloidin-stabilized actin filaments. Fluorescence intensity along five filaments was measured for five frames. Data are expressed as the mean ± SD of 25 measurements. **(D)** Cosedimentation of GFP-Rng2CHD with 3 µM of actin filaments. Similar molar amounts of actin and GFP-Rng2CHD were recovered in the pellet fractions when [GFP-Rng2CHD] was 5 and 6 µM. The GFP-Rng2CHD preparation was contaminated by a bacterial ∼30 kDa protein.

To directly confirm that sparsely bound Rng2CHD potently inhibits actomyosin movements on muscle HMM, we prepared Rng2CHD fused with GFP to its N-terminus via a 16-residue linker (GFP-Rng2CHD), and determined the binding ratio of GFP-Rng2CHD to actin protomers from the fluorescence intensity of GFP. GFP-Rng2CHD strongly inhibited actin filament movement on muscle HMM in a manner similar to Rng2CHD (Figure 3A). We then measured the fluorescence intensity of bound GFP-Rng2CHD per unit length of actin filaments. Fluorescence intensity increased depending on the GFP-Rng2CHD concentration in the buffer, and was saturated in the presence of 5 µM GFP-Rng2CHD (Figure 3B, 3C). The co-sedimentation assay showed that one molecule of GFP-Rng2CHD binds to one molecule of an actin protomer in the presence of 5 µM GFP-Rng2CHD (Figure 3D). Therefore, we regarded this saturated fluorescence intensity as a one-to-one binding state, and used it as the reference to calculate how much GFP-Rng2CHD binds to actin protomers when movements were inhibited in the presence of lower concentrations of GFP-Rng2CHD. This fluorescence-based direct quantification also demonstrated that sparsely bound GFP-Rng2CHD strongly inhibits actin movements on muscle HMM (Table 1), although based on these directly measured binding ratios of GFP-Rng2CHD to actin protomers, a higher binding ratio was needed to obtain the same degree of motility inhibition than that estimated from *K_d_* using unlabeled Rng2CHD. Similarly higher binding ratios needed to obtain the same degree of motility inhibition were obtained when the binding ratio of GFP-Rng2CHD was estimated from *K_d_* that was derived from the cosedimentation experiments (Supplementary Information 1).

### Rng2CHD cooperatively changes the structure of actin filaments, accompanying supertwisting and local kinks or structural distortions

Based on the very low binding ratio of Rng2CHD required to inhibit movement of actin filaments on muscle myosin II, we inferred that sparsely bound Rng2CHD somehow induces global structural changes in actin filaments and inhibits actomyosin II movement. To examine whether sparsely bound Rng2CHD actually changes the structure of actin filaments, we observed actin filaments in the presence of Rng2CHD using negative stain electron microscopy and high speed atomic force microscopy (HS-AFM).

For HS-AFM observations, actin filaments were loosely immobilized on a positively charged lipid bilayer, the condition which we previously used to detect cofilin-induced supertwisting of actin filaments (Ngo et al., 2015). Periodic patterns representing half helices of the double-helical structures were clearly observed, in which the tallest parts in each half helix, or the crossover points, are shown in a brighter color (Figure 4A). In the presence of 0.59 µM actin and 20 nM Rng2CHD, no bound Rng2CHD molecules were detected, although the binding ratio estimated from *K_d_* was 1.3%. When the concentration of Rng2CHD was increased to 0.25 µM, which corresponds to the estimated binding ratio of 15%, we were able to detect sparse and transient binding events (Figure 4B and Video 6). In the image shown in Figure 4B, there are approximately 57 half helices, or approximately 740 actin protomers, implying that there must be 110 bound Rng2CHD molecules in this image. However, there is only one Rng2CHD-derived bright spot in this image. Those bright spots appear and disappear transiently, but the number of Rng2CHD spots that could be detected simultaneously in this imaging field did not exceed two during the 14.5 s of observation (Video 6). Regarding this apparently large discrepancy, it is notable that all the Rng2CHD-derived bright spots appeared at the crossover points of the double helix (Video 6). Moreover, all the Rng2CHD-derived bright spots were as large as or larger than actin protomers (42 kDa) and some bright spots obviously consisted of two smaller bright spots. We thus speculate that what we imaged were clusters of two or more Rng2CHD molecules bound near the crossover points, and individual bound Rng2CHD molecules and the clusters that bound along the filament sides were not efficiently imaged due to the small size (21 kDa).

**Figure 4.**
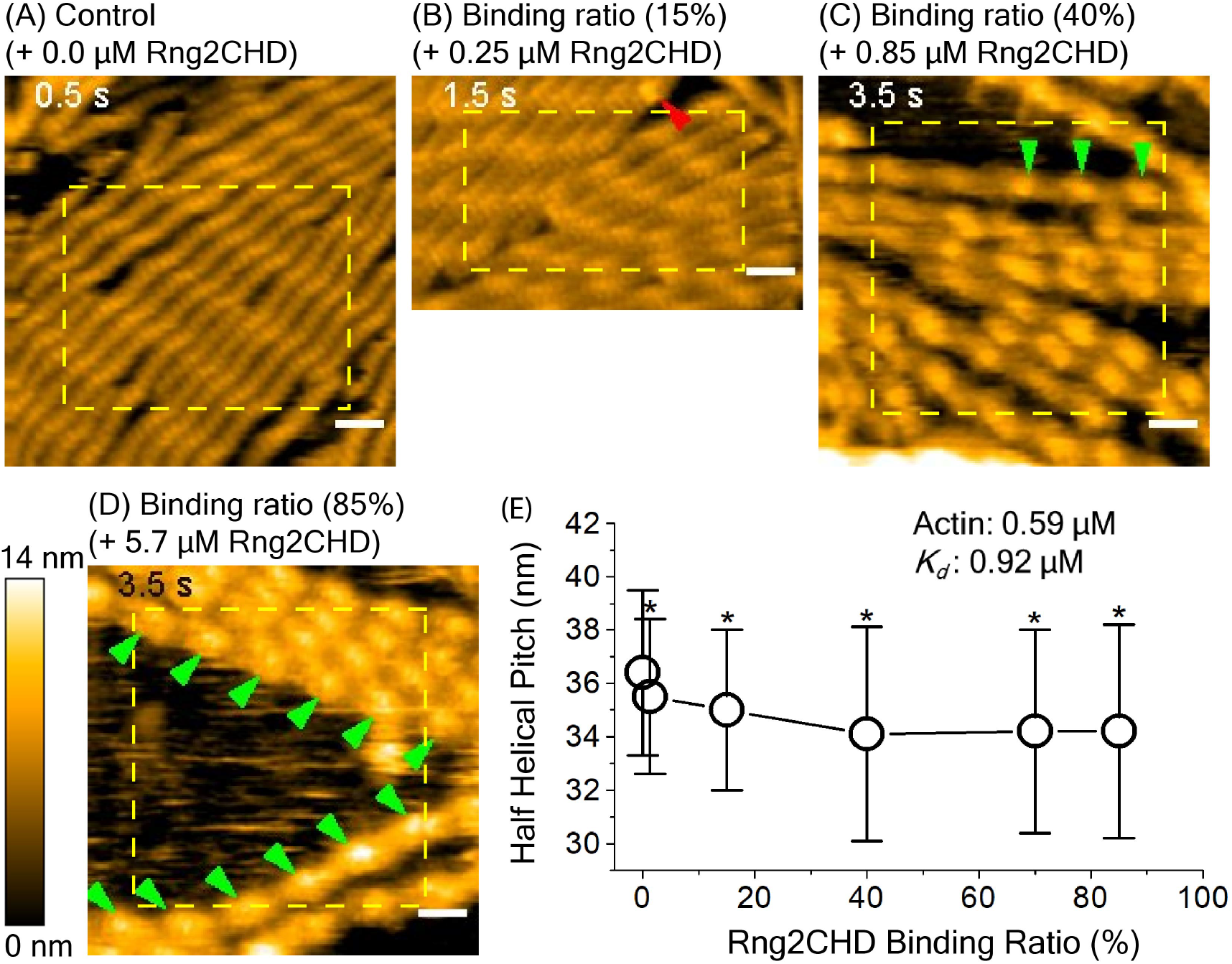
HS-AFM imaging and analysis of half helical pitch (HHP) of actin filaments at different Rng2CHD binding ratios. Actin filaments were premixed with Rng2CHD at different concentrations in a tube for 10 min at RT to achieve equilibrium binding. These protein mixtures (68 µl) were introduced into an observation chamber for HS-AFM imaging, and actin filaments with and without bound Rng2CHD were gently immobilized onto positively-charged lipid bilayer formed on mica. In all experiments, the concentration of actin filaments was fixed at 0.59 µM while the concentrations of Rng2CHD were varied at 0, 0.020, 0.25, 0.85, 2.6, and 5.7 µM. Typical images of actin filaments at different Rng2CHD binding ratios are shown in **A-D**. Red and green arrowheads denote isolated and series of Rng2CHD clusters, respectively. Note that only selected series of the Rng2CHD clusters are marked. Bars: 25 nm. Using those images, half helical pitches (HHPs) were estimated by measuring the distances between the peaks of two neighboring two half helices. Half helices in the yellow rectangles were subjected to the HHP measurements, regardless of the presence or absence of Rng2CHD clusters. The Rng2CHD binding ratio was estimated by using the *K_d_* value of 0.92 µM. The correlation between Rng2CHD binding ratio and HHP of actin filaments is shown in **E**. The values are mean HHP ± SD. Note that the position of the peak (highest point of the actin protomer that is closest to the crossover point) does not necessarily coincide with the crossover point of two helices that connect the centers of the mass of actin protomers in each protofilament, and there can be up to 5.5/2 nm error between the two positions. This error would not affect the mean of HHPs, since they would be averaged out when multiple HHPs are measured consecutively along actin filaments, but would contribute to the larger SD values (Ngo et al., 2015). The statistical differences of the mean HHP of control actin filaments (0 µM Rng2CHD) and that at different Rng2CHD binding ratios (*, *p* ≦ 0.001, two independent populations *t*-test) were calculated. **<u>Related to Video 6 and Table Supplement 1</u>.**

In the presence of 0.85 and 2.6 µM Rng2CHD, which correspond to the estimated binding densities of 40% and 70%, respectively, progressively larger fraction of crossover points became brighter, while other crossover points remained unchanged (Figure 4C), again suggesting the propensity of Rng2CHD to form clusters. In the presence of 5.7 µM Rng2CHD, which corresponds to the estimated binding ratio of 85%, most of the crossover points were brighter (Figure 4D), implying a nearly saturated binding. We then measured half helical pitches (HHP) of actin filaments by measuring the distances between the crossover points under equilibrium binding conditions, and found that Rng2CHD induced shortening of HHP, or supertwisting (Figure 4E). Strikingly, the supertwisting conformational changes nearly saturated at 0.85 µM Rng2CHD, when the estimated binding ratio was 40%. These results clearly demonstrate that sparsely bound Rng2CHD induces cooperative conformational changes to actin filaments. Another notable feature of actin filaments in the presence of Rng2CHD is the transient local untwisting of the helix to result in two separate protofilaments (Figure Supplement 3 and Video 7).

We also employed frequency modulation atomic force microscopy (FM-AFM) to observe the structural changes on actin filaments induced by Rng2CHD (Figure Supplement 4A). FM-AFM has emerged as a powerful tool to provide nanostructural information for various surfaces/interfaces and biological samples in liquid environments with unprecedented spatial resolution (Giessibl, 2003; Ido et al., 2013). FM-AFM observation confirmed Rng2CHD-induced decrease of HHP (Figure Supplement 4B).

Negative staining and electron microscopic observation of actin filaments in the absence of Rng2CHD showed long and straight filaments (Figure 5A), and the higher magnification images were consistent with previously known helical structures (Figure 5E). However, when the actin filaments were allowed to interact with Rng2CHD in solution, then deposited onto carbon-coated grids and negatively stained (Figure 5B-5D), the appearance of filaments became irregular and more filaments became bundled. At higher concentrations (200 nM – 1 µM) of Rng2CHD, the filaments were frequently distorted, kinked, or even fragmented. Even those filaments that appeared straight often showed irregular helical structures at high magnifications (Figure 5F-5H): the two actin protofilaments appeared to be separated in some portions of the filaments, so that a dark straight line was observed in between two parallel protofilaments (indicated by yellow brackets). The observed separation of the protofilaments presumably correspond to the parallel protofilaments observed by HS-AFM (Figure Supplement 3), indicating that they are not artifacts of HS-AFM or negative staining. Such Rng2CHD-induced local untwisting conformational changes appear at odds with the Rng2CHD-induced supertwisting observed by HS-AFM. Since the local untwisting was frequently observed in the medium concentration range of Rng2CHD, we speculate that the local untwisting is a compensatory conformational change that is induced by Rng2CHD-induced local supertwisting and facilitated by Rng2CHD-induced weakening of interactions between the protofilaments.

**Figure 5.**
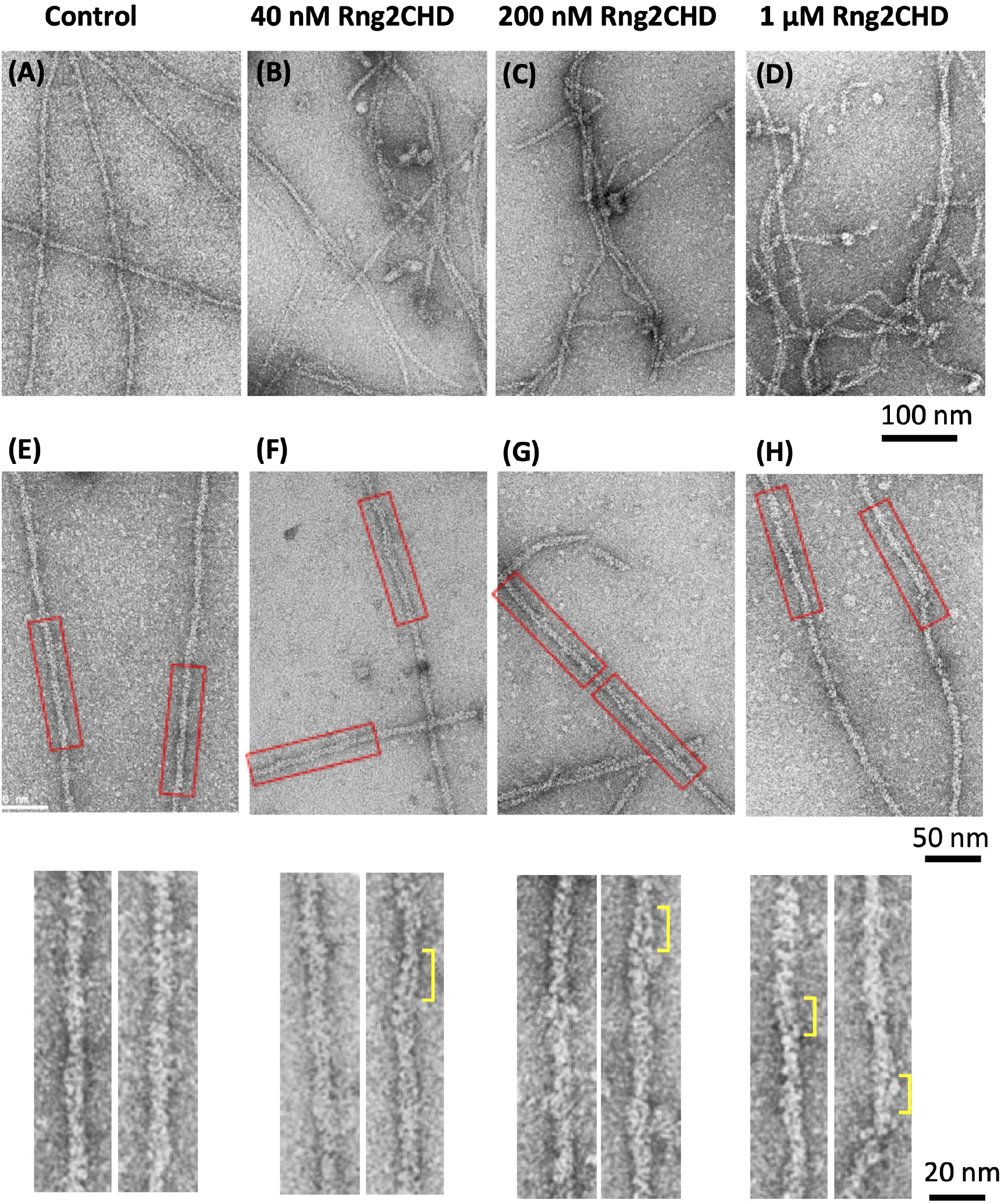
Rng2CHD deforms the helical structures of actin filaments. Electron micrographs of negatively stained actin filaments in the absence **(A, E)** and presence of 40 nM **(B, F)**, 200 nM **(C, G)** and 1 µM **(D, H)** Rng2CHD. In the presence of Rng2CHD, the filaments were often bundled and kinked. In the presence of high concentrations of Rng2CHD, the filaments were highly deformed and showed discontinuities (**C** and **D**), which can be explained by altered interactions between neighboring actin protomers. **E**, **F**, **G** and **H** are higher magnifications of the straight portions of the filaments under each condition, and the boxed regions are further magnified in the bottom row. In some parts, the two actin protofilaments look separated (yellow brackets), suggesting that Rng2CHD somehow reduces, or at least changes, the interaction between the two protofilaments at sub-stoichiometric binding ratios. In the absence of Rng2CHD, such irregular filament structures were not observed **(A, E)**.

### Steady state actin-activated muscle S1 ATPase is only weakly inhibited by Rng2CHD

To gain insight into the mechanism by which structural changes of actin filaments induced by Rng2CHD inhibit motility by muscle myosin II, we investigated the effects of Rng2CHD on actin-activated ATPase activity of muscle myosin II subfragment-1 (S1). Actin-activated S1 ATPase was moderately inhibited (approximately 50%) by the highest concentration of Rng2CHD tested (5 μM; Figure 6). In the presence of 0.33 μM and 0.82 μM Rng2CHD and 24 μM of actin filaments, the actin-activated S1 ATPase activity was not inhibited in a statistically significant manner. Under those conditions, the binding ratios of Rng2CHD to actin were calculated as 1.3% and 3.7% (Eq. 2 in Materials and Methods), which caused a 50% and 82% reduction in the speed of actin movement, respectively, by muscle HMM (Figure 1A and Table 1). In the presence of 1.9 µM Rng2CHD, S1 ATPase was inhibited by 29%, which was statistically significant (p<0.03). In this condition, the binding ratio was calculated as 7.6%, which inhibited 96% of sliding speed. Thus, the inhibition of actin-activated S1 ATPase activity was much weaker and disproportional to the inhibition of movement (Figure 6). This indicates that Rng2CHD-induced strong inhibition and stalled actin movements on muscle HMM do not necessarily accompany inhibition of the ATPase cycle.

**Figure 6.**
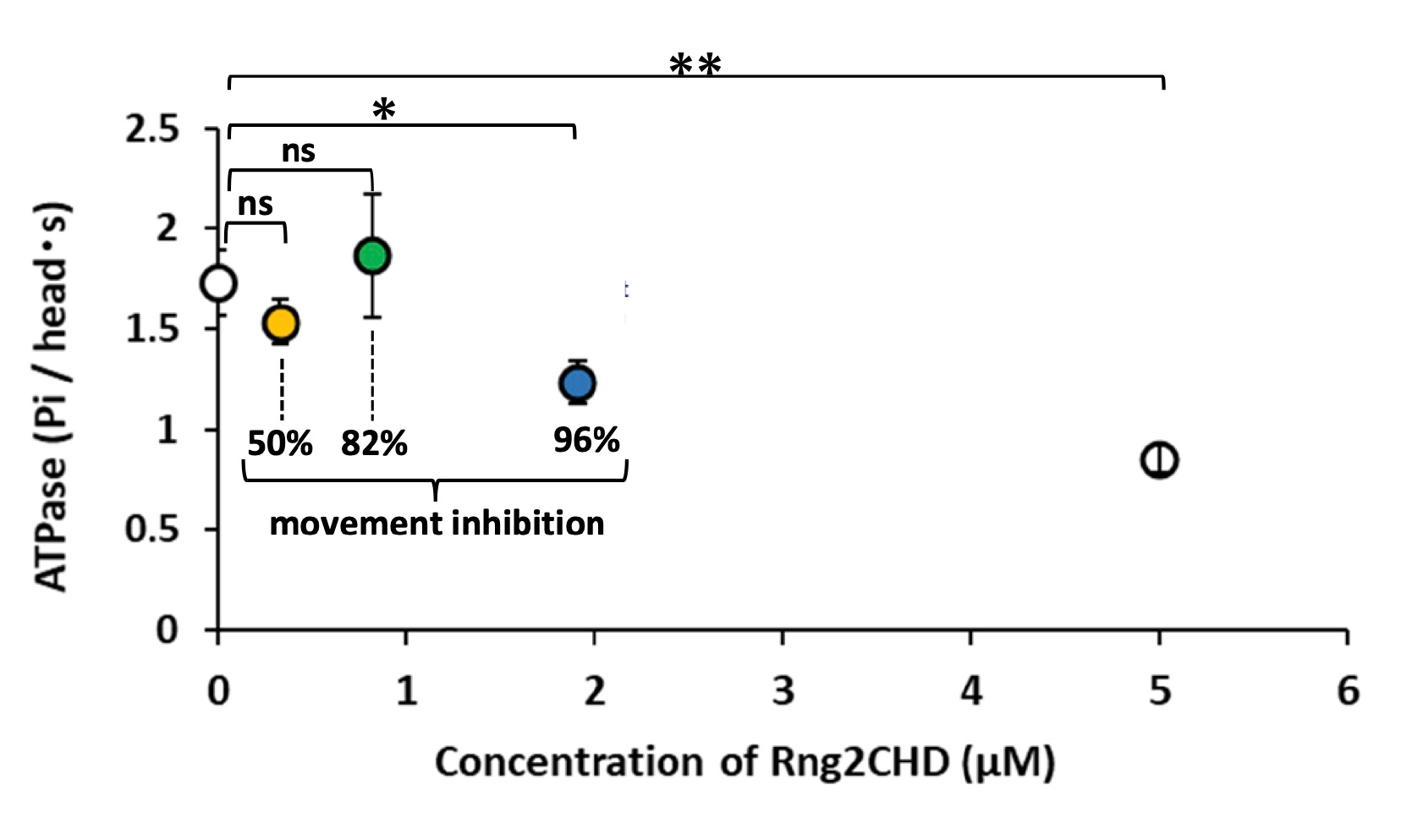
Actin-activated muscle S1 ATPase in the presence of Rng2CHD. The orange (0.33 µM Rng2CHD), green (0.82 µM Rng2CHD) and blue (1.9 µM Rng2CHD) plots were measured in the presence of Rng2CHD concentrations that were expected to bind to actin protomers at binding ratios which caused a 50%, 82% and 96% reduction of actomyosin movement speed on muscle HMM, respectively during *in vitro* motility assays. Note that the concentration of Rng2CHD to achieve the same binding ratio is very different between this ATPase experiment and the motility assays because the concentration of actin is very different between the two experiments. Data are expressed as the mean ± SD of three independent experiments. “ns” indicates that the differences are not statistically significant; “*” and “**” indicate statistically significant differences with a *p* value < 0.03 and < 0.002, respectively, according to a Student’s *t*-test.

### Rng2CHD inhibits the steady-state binding of muscle S1 to actin filaments in the presence of ADP, but not in the presence of ATP

We examined the possibility that Rng2CHD might affect the affinity between actin filaments and myosin motor when it inhibits motility by muscle HMM. First, we performed a co-sedimentation assay of actin filaments and muscle S1, and found that Rng2CHD did not significantly inhibit steady-state binding of S1 to actin filaments in the presence of ATP (Figure 7A, 7C). However, Rng2CHD weakly but statistically significantly inhibited the binding of S1•ADP to actin filaments in the presence of ADP in the buffer (Figure 7B, 7C).

**Figure 7.**
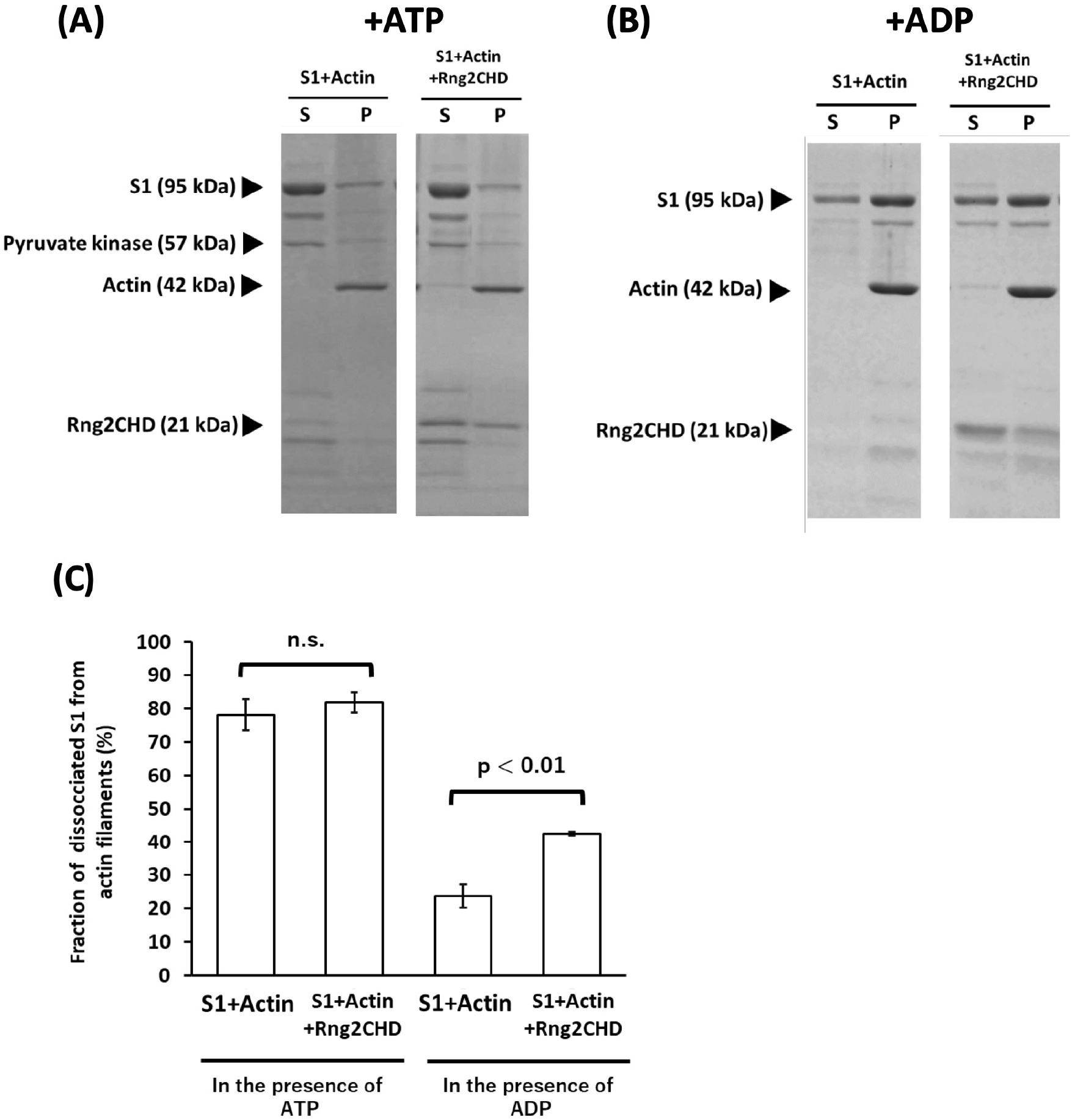
Rng2CHD inhibits the steady-state binding of S1 to actin filaments in the presence of ADP, but not in the presence of ATP. **(A, B)** Co-sedimentation assay of S1 and actin filaments in the presence of Rng2CHD and 2 mM ATP (A) or 2 mM ADP (B). **(C)** Fraction of S1 dissociated from actin filaments was compared with and without Rng2CHD. Rng2CHD significantly increased the fraction of dissociated S1 from actin filaments in the presence of ADP (Student’s *t*-test, *p*<0.01), but not in the presence of ATP. Data are expressed as the mean ± SD of three independent experiments.

A co-sedimentation assay using muscle myosin II filaments showed that Rng2CHD only weakly bound to myosin II under the conditions employed in the *in vitro* motility assays (Figure Supplement 5).

We also employed HS-AFM to directly observe transient binding of muscle S1 molecules to actin filaments in the presence of ATP. At a scan speed of 0.5 s per field of view, transient binding of S1 to actin filaments was rarely observed in the presence of 500 μM ATP alone, but was frequently observed in the presence of 50 μM ATP and 1 mM ADP. S1 molecules were easily identified based on their size and shape, whereas individual bound Rng2CHD molecules were not visualized as described earlier. We analyzed images scanned between 1 and 2 min after the addition of S1, and visually counted the number of transient binding events of S1 molecules (Figure 8A). The binding dwell time of S1 molecules on the top of the filament was shorter than those bound along the sides of the filaments. Therefore, we separately counted the S1 molecules bound on the top and along the sides of the filaments. The number of S1 molecules that transiently bound to actin filaments was significantly lower when 12 nM Rng2CHD, the concentration that caused 50% inhibition of motility on muscle HMM, was added before the addition of S1 (Figure 8B, 8C). It was thus directly confirmed that sparsely bound Rng2CHD affected the binding of S1 to actin filaments in the presence of ATP and ADP.

**Figure 8.**
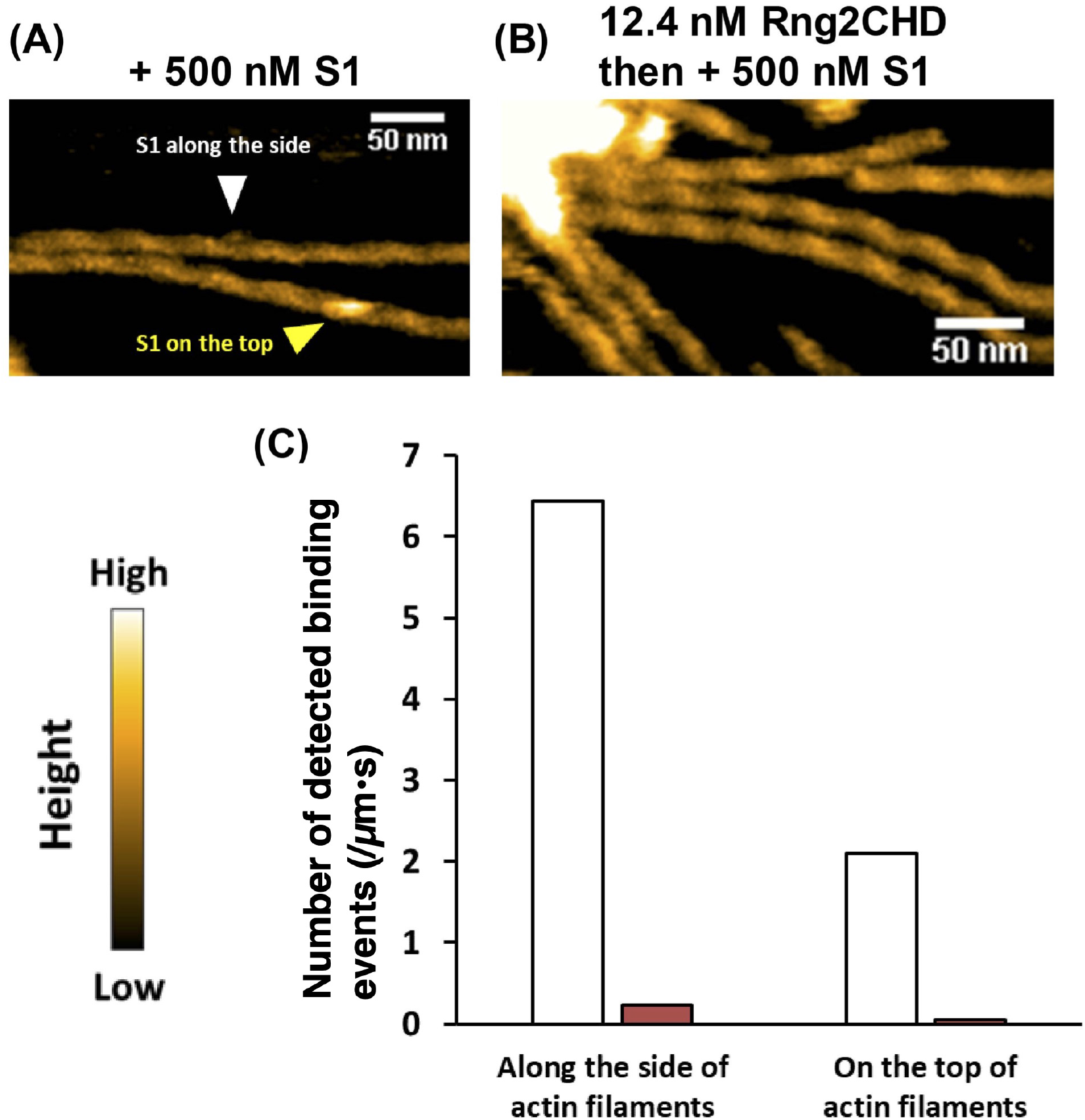
Rng2CHD significantly decreases the number of S1 molecules bound along actin filaments in the presence of ATP and ADP. **(A, B)** HS-AFM images of actin filaments interacting with S1 in the presence of 50 µM ATP and 1 mM ADP. The images were scanned at about 2 min after the addition of 500 nM S1. S1 molecules that bound on the top and along the side of the filaments are indicated by yellow and white arrowheads, respectively. In **(A)**, 500 nM S1 was added to actin filaments. In **(B)**, in contrast, actin filaments were preincubated with 12 nM Rng2CHD for 15 min, then 500 nM S1 was added. **(C)** Number of observed S1 binding events on the top or along the sides of the filaments in the presence of 50 μM ATP and 1 mM ADP. The values were normalized by the total length of the measured filaments and time. White bars: 500 nM S1 was added to actin filaments. Red bars: Actin filaments were preincubated with Rng2CHD, and then 500 nM S1 was added. The number of bound S1 molecules was counted in the images scanned between 1 and 2 min after the addition of S1.

The result that transient binding of S1 molecules to actin filaments was hardly observed in the presence of 500 μM ATP alone suggests that, in the presence of 50 µM ATP and 1 mM ADP, HS-AFM presumably detected S1•ADP bound to actin filaments before the low concentration of ATP in the presence of excess ADP slowly disrupted the binding. The decrease in the number of detected S1 molecules caused by Rng2CHD can be interpreted in the following two ways: (1) the number of transient binding events of S1 decreased, or (2) the duration of each binding event was shortened. Hypothesis (1) predicts that actin-activated S1 ATPase is also very strongly inhibited by Rng2CHD, which was not the case. We thus concluded that the unstable binding of S1•ADP to actin filaments caused by Rng2CHD shortened the duration of transient binding of S1•ADP to actin filaments, and decreased the efficiency of detection of transient binding by HS-AFM.

### Rng2CHD decreases the fluorescence of HMM-GFP along actin filaments in the presence of ATP

We previously reported that when HMM of *Dictyostelium* myosin II fused with GFP was allowed to interact with actin filaments in the presence of a very low concentration of ATP, HMM-GFP formed clusters along actin filaments (Tokuraku et al., 2009; Hirakawa et al., 2017). This was interpreted to represent local polymorphism of actin filaments, such that some segments of the filaments have a higher affinity for HMM than other parts of the filament, and HMM-GFP preferentially repeats transient binding to those segments. In the presence of a high concentration of ATP, no fluorescent clusters were observed, and in the absence of ATP, HMM-GFP uniformly bound along the entire filaments (Tokuraku et al., 2009).

We speculated that, in order for the detectable fluorescent clusters to form, HMM needs to bind to actin filaments repetitively and transiently, although the dwell time of each binding event must be long enough to allow visualization. Based on this hypothesis, we employed fluorescence microscopy to observe how Rng2CHD affected the formation of *Dictyostelium* HMM-GFP clusters along actin filaments in the presence of a very low concentration of ATP. Numerous fluorescent spots, each representing an HMM-GFP cluster, were observed along actin filaments in the presence of 0.5 µM ATP and in the absence of Rng2CHD. In the presence of 1 nM Rng2CHD, the number and fluorescence intensity of fluorescent spots were significantly reduced, and fluorescent spots were apparently absent in the presence of 10 nM Rng2CHD (Figure 9A, 9C). In contrast, Rng2CHD did not affect the binding of HMM-GFP to actin filaments in the nucleotide-free state (Figure 9B, 9C).

**Figure 9.**
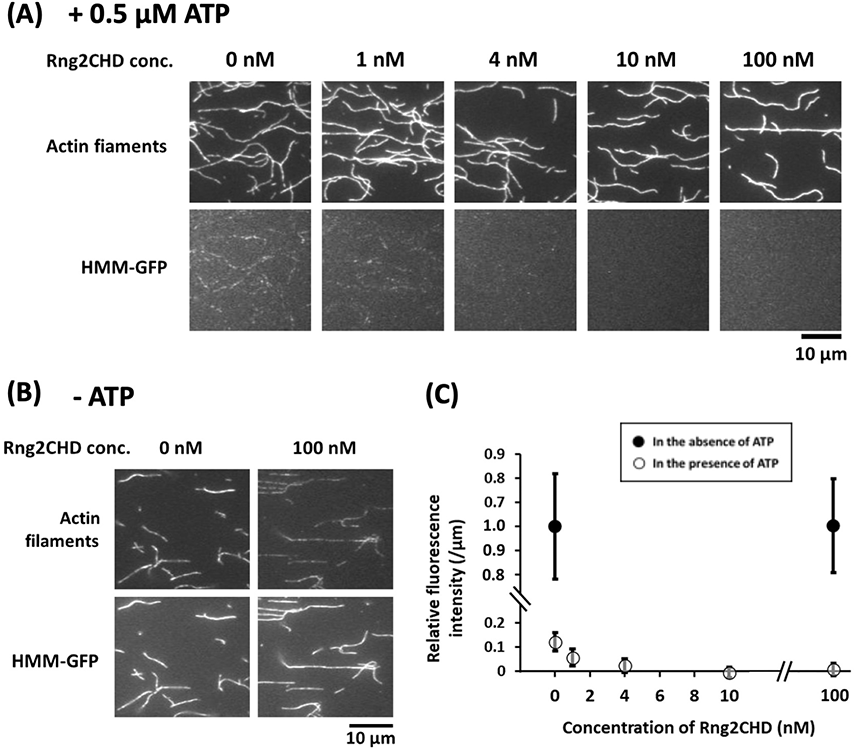
Rng2CHD decreases the fluorescence of HMM-GFP along actin filaments in the presence of ATP. Fluorescence micrographs of actin filaments labeled lightly with Alexa647 phalloidin and *Dictyostelium* HMM-GFP in the presence of 0.5 µM ATP **(A)** and in the nucleotide-free state **(B)**. In each panel, the top row shows the fluorescence of Alexa647, and the bottom row shows that of GFP. Fluorescence intensity of GFP was measured along each actin filament and divided by the length of the filament, and mean and SD of the resultant values were calculated for each condition (N=20). These normalized GFP fluorescence intensities were further normalized against the value in the absence of ATP and Rng2CHD to obtain relative fluorescence intensity (**C**). Filled circles show the data in the absence of ATP and the open circles show the data in the presence of 0.5 µM ATP. The binding ratio of Rng2CHD along the actin filaments, estimated from *K_d_*, are 0.1, 0.4, 1.1 and 9.8% for the Rng2CHD concentrations of 1.0, 4.0, 10 and 100 nM, respectively.

## Discussion

### Structural changes of actin filaments induced by sparsely bound Rng2CHD inhibit actomyosin II movement

Rng2CHD, the actin-binding domain of Rng2, strongly inhibited actomyosin II motility, and particularly potently on skeletal muscle myosin II. Inhibition occurred along the entire length of the filament on muscle HMM, when actin filaments were only sparsely decorated by Rng2CHD or GFP-Rng2CHD (Figure 1A, 3A, Video 2, and Table 1). Since binding of Rng2CHD or GFP-Rng2CHD was sparse, the inhibition of motility could not be due to steric hindrance or direct competition for a binding site on actin molecules. We thus inferred that sparsely bound Rng2CHD induced some cooperative structural changes in actin filaments, and these inhibited the productive interaction between actin filaments and myosin II. Previous studies advocated that structural changes of actin filaments modulate affinities for various ABPs (Ngo et al., 2016; Shibata et al., 2016; Harris et al., 2020). For example, cofilin cooperatively binds to actin filaments to form clusters along the filaments, reducing the helical pitch of filaments in the cluster by 25% (McGough et al., 1997). Notably, this structural change was propagated to the neighboring cofilin-unbound bare region (Galkin et al., 2001; Ngo et al., 2015), and this was accompanied by decreased affinity for muscle S1 in the presence of ATP (Ngo et al., 2016). In this study, HS-AFM observation showed that Rng2CHD cooperatively changed the structure of actin filaments accompanying supertwisting even when Rng2CHD was only sparsely bound to actin filaments (Figure 4E). Discontinuities and kinks of the actin filaments, as well as the dark straight lines between the two protofilaments observed by electron microscopy (Figure 5), suggest that the interaction between actin protomers were altered by Rng2CHD at sub-stoichiometric binding densities. Although the extent of supertwisting by Rng2CHD (∼5%) was much smaller than that caused by cofilin, it is notable that Rng2CHD and cofilin share two properties, namely supertwisting of the actin helix, and a decreased affinity for myosin II. Elucidating whether there is a causal relationship between the two properties or they are mere coincidence needs further investigations.

We consider two possible mechanisms by which sparsely bound Rng2CHD inhibits actomyosin II movements. The first mechanism proposes that one or two actin protomers in direct contact with the bound Rng2CHD molecule undergo structural changes, and those affected actin protomers bind persistently to myosin II motors even in the presence of ATP, acting as a potent break. The second mechanism assumes that a bound Rng2CHD molecule changes the structure of multiple actin protomers, and the affected actin protomers become unable to productively interact with myosin II. The first mechanism predicts that Rng2CHD should increase the amount of co-sedimented S1 in the presence of ATP, which was not the case (Figure 7A, 7C). Direct observation of actin binding of *Dictyostelium* myosin II HMM-GFP in the presence of a very low concentration of ATP also demonstrated that Rng2CHD decreased the affinity between actin filaments and myosin II motors in a concentration-dependent manner (Figure 9A, 9C). Furthermore, buckling of the moving actin filaments on muscle HMM-coated surfaces, indicative of local inhibition of the movement, was rarely observed in the presence of various concentrations of Rng2CHD (Figure 1C). The tendency of actin filaments on *Dictyostelium* myosin II to slide sideways and to detach from the myosin-coated surface is also inconsistent with the local break hypothesis. Those reasons led us to reject the first mechanism and conclude that force generation by myosin II is inhibited in broad sections of actin filaments that are not in direct contact with Rng2CHD.

Hereafter, we resolve the inhibition process into two aspects, and discuss their respective mechanisms. The first is the mechanism by which sparsely bound Rng2CHD causes global structural changes in actin filaments. The second is the mechanism by which the structural changes of actin filaments inhibit actin motility driven by myosin II.

### The mechanism by which Rng2CHD causes global structural changes in actin filaments

We now consider two hypotheses for the mechanism by which one molecule of Rng2CHD changes the structure of multiple actin protomers. The first is a cooperative structural change, in which one molecule of Rng2CHD bound to one actin protomer changes the structure of multiple neighboring actin protomers in the same filament. Such cooperative propagation of conformational changes has been reported for many ABPs. The best characterized case is the propagation of a supertwisted structure in the cofilin clusters to neighboring bare zones (Galkin et al., 2001; Ngo et al., 2015). Moreover, a single molecule of gelsolin bound to the barbed end of an actin filament changes the structure of all actin protomers in the filament (Orlova et al., 1995). Other ABPs such as myosin II (Oosawa et al., 1973; Miki et al., 1982; Prochniewicz et al., 2010), tropomyosin (Khaitlina et al., 2017), α-actinin (Singh et al., 1981) and formin (Papp et al., 2006) also cause cooperative structural changes of actin filaments even at significantly sub-stoichiometric concentrations to actin monomers. The second hypothesis is the memory effect, in which Rng2CHD molecules repeat transient binding to different actin protomers, which remain in an altered conformation for a certain period of time after dissociation of Rng2CHD. If there is a memory effect, Rng2CHD can alter the structure of entire actin filaments even when its binding ratio to actin protomers is low. We speculate that either or both of these two mechanisms, cooperative structural change and memory effect, underlie the global structural changes in actin filaments induced by sparsely bound Rng2CHD.

### The mechanism by which structural changes of actin filaments inhibit actomyosin II motility

Rng2CHD inhibited steady-state binding of S1•ADP to actin filaments (Figure 7B, 7C). Consistent with this result, HS-AFM demonstrated that Rng2CHD significantly reduced the binding dwell time of muscle S1 molecules on actin filaments in the presence of 50 µM ATP and 1 mM ADP (Figure 8). Moreover, fluorescence microscopy demonstrated that Rng2CHD significantly decreased the region along actin filaments where *Dictyostelium* HMM-GFP fluorescence was observed in the presence of 0.5 μM ATP (Figure 9A).

Based on this conclusion, we propose two possible mechanisms for inhibition of actomyosin II movement caused by Rng2CHD, in the framework of the swinging lever arm model (Huxley, 1969; Cooke et al., 1984; Tokunaga, 1991; Uyeda et al., 1996) tightly coupled with the actomyosin ATPase cycle (Lymn and Taylor, 1971). The first mechanism proposes that phosphate release from myosin II•ADP•Pi is promoted normally by actin filaments that have been structurally altered by Rng2CHD, but without the lever arm swing that normally accompany the phosphate release. Consequently, myosin II•ADP, which does not have the authentic post-power stroke structure, cannot gain the normal high affinity to actin filaments. The second mechanism assumes that although the lever arm swing occurs following phosphate release, the myosin II motor domain•ADP slips at the contact surface with actin filaments, or myosin II•ADP dissociates from actin filaments, because of the low affinity between myosin II•ADP and the structurally altered actin filaments. This would lead to a failure of myosin II•ADP to maintain the tension, generated by the swing of the lever arm, to drive the movement of the actin filaments. The two inhibition mechanisms are both derived from a defective interaction between the affected actin and myosin motor carrying ADP, and may not be mutually exclusive. The two myosin IIs used in this study, i.e., skeletal muscle myosin II and *Dictyostelium* myosin II, appeared to respond differently to Rng2CHD-affected actin filaments, but the differences can be explained, at least in part, by known quantitative differences between the two myosin IIs within the framework of this proposed mechanism of inhibition (Supplementary Information 2).

Actin movements by myosin V HMM were even more different in terms of sensitivity to Rng2CHD, in that the sliding velocity by myosin V was not appreciably affected by up to 1 µM of Rng2CHD. In line with this finding, it is worth mentioning that two actin mutations, M47A (Kubota et al., 2009) and G146V (Noguchi et al., 2012), inhibit actin movements on muscle myosin II, but not on myosin V. Further studies are needed to understand the mechanism by which myosin II and V respond qualitatively differently to inhibition by Rng2CHD and certain actin mutations.

### Future studies

Further investigations are also warranted to reveal the structural aspects of the inhibition of actomyosin II movement induced by Rng2CHD, and to clarify the relationship between the structural changes of actin filaments and actomyosin II motility.

Phosphorylation of the myosin light chain (Higashi-Fujime, 1983; Sellers et al., 1985; Griffith et al., 1987) and calcium regulation via tropomyosin and troponin (Ebashi and Kodama, 1965; Ebashi and Kodama, 1966) are two widely known major regulatory mechanisms of actomyosin II movements. Additionally, it has been reported that caldesmon and calponin inhibit the movement of actin filaments on smooth muscle myosin II (Shirinsky et al., 1992). Of those two classic regulators of smooth muscle contraction, calponin is homologous to Rng2CHD. Moreover, these ABPs are similar to Rng2CHD in that they inhibit actomyosin II movements even with sparse binding to actin filaments (Shirinsky et al., 1992), although the cooperativity of motility inhibition on muscle HMM is weaker than that of Rng2CHD. More information is needed to further discuss the mechanistic similarities and differences among the inhibition by Rng2CHD, calponin and caldesmon.

The physiological significance of the inhibitory effect of Rng2CHD on actomyosin II is another unresolved issue. Since contraction of the CR appears to be regulated in an inhibitory manner (Supplementary Information 3), it is plausible that Rng2CHD plays a role in this regulatory process. Recently, Palani et al. (2021) demonstrated that Rng2CHD, or “curly” according to their nomenclature, that was loosely immobilized on a lipid membrane formed rings of actin filaments *in vitro*, suggesting that it is involved in the formation of CRs *in vivo*. However, a previous truncation study showed that *S. pombe* cells expressing mutant Rng2 lacking the CHD are able to assemble and contract CRs normally (Tebbs and Pollard, 2013). Molecular and cell biological studies are thus needed to understand the physiological role of Rng2CHD, including possible functional redundancy with α-actinin, overexpression of which significantly slows cytokinesis in mammalian cells (Mukhina et al., 2007).

## Materials and Methods

### Protein purification

Actin was purified from rabbit skeletal muscle acetone powder (Spudich and Watt, 1971; Pardee and Spudich, 1982). HMM and S1 of muscle myosin II were prepared by digestion of rabbit skeletal muscle myosin with papain and *α*-chymotrypsin, respectively (Margossian and Lowey, 1982). *Dictyostelium* full length myosin II and HMM-GFP were purified as described previously (Ruppel et al., 1994; Tokuraku et al., 2009). The HMM version of human myosin V with a FLAG-tag at the N-terminus and a c-myc tag at the C-terminus was coexpressed with calmodulin in insect cells and purified using a method described previously (Watanabe et al., 2006).

In our previous study, we used Rng2CHD fused with a His-tag at the N-terminus, and reported that His-Rng2CHD bundles actin filaments (Takaine et al., 2009). However, we subsequently discovered that the His tag enhances the affinity of Rng2CHD for actin filaments, and untagged Rng2CHD has very poor actin bundling activity while retaining actin binding activity (Figure Supplement 6). In this study, therefore, we used untagged Rng2CHD prepared as follows. The gene encoding Rng2CHD (Takaine et al., 2009) was inserted at the *Hind*III and *Pst*I sites of the pCold-TEV vector (Ngo et al., 2015), which had a TEV protease recognition sequence between the 6×His sequence and the multiple cloning site of pColdI (Takara Bio, Kusatsu, Japan). The amino acid sequence of His-TEV Rng2CHD was MNHKVHHHHHHIEGRHM<u>ENLYFQG</u>TLEGSEFKLDVNVGL…(Rng2CHD)…LPNFKA, where the underline shows the TEV recognition sequence.

Rng2CHD was expressed in BL21 *Escherichia coli* (Takara Bio) according to the instructions provided by the manufacturer of pColdI. The cells were lysed by sonication in 2 mM 2-mercaptoethanol, 0.3% Triton X-100, 0.1 mM phenylmethylsulfonylfluoride, 400 mM NaCl, 10 mM imidazole (pH 7.4) and 20 mM Hepes (pH 7.4) on ice. The homogenate was clarified by centrifugation and mixed with Ni Sepharose 6 Fast Flow (GE Healthcare, Chicago, IL). After extensive washing, the peak fractions eluted by 7 mM 2-mercaptoethanol, 400 mM imidazole (pH 7.4) and 10 mM Hepes (pH 7.4) were combined and supplemented with His-tagged TEV protease at a 1/10 molar amount of proteins to separate the His-tag from Rng2CHD at the cleavage site for TEV protease. After dialysis against 50 mM KCl, 0.1 mM DTT and 10 mM Hepes (pH 7.4) overnight at 4°C, the protein solution was clarified by centrifugation and passed through Ni Sepharose 6 Fast Flow in a column to remove the released His-tag, His-TEV protease and uncleaved His-tagged Rng2CHD. This was followed by concentration with a centrifugal concentrator (Amicon Ultra-15 3 k device, Merck Millipore, Burlington, MA), and after supplementing with 10 mM DTT, aliquots were snap-frozen in liquid nitrogen and stored at −80°C.

To fuse GFP to the N-terminus of Rng2CHD, the GFP gene with a S65T mutation was inserted at the *Kpn*I and *BamH*I sites of pCold-TEV-Rng2CHD. A Gly-based 16 amino acid-residue linker sequence was inserted between the GFP gene and the Rng2CHD gene, so that the expressed GFP would not spatially inhibit the binding of Rng2CHD to the actin filament. The resultant amino acid sequence of GFP-Rng2CHD was MNHKVHHHHHHIEGRHM<u>ENLYFQG</u>TMSKGE…(GFP)…MDELY<u>GGSEFGSSGSSGSSKL</u>DV NVGL…(Rng2CHD)…LPNFKA, where the underline shows the TEV recognition sequence and the double underline shows the linker sequence. GFP-Rng2CHD was expressed in Rosetta DE3 *E. coli* (Merck Millipore), and purified basically in the same way as Rng2CHD, except that the protein was further purified by anion exchange chromatography with an Econo-pac High Q cartridge (BIO-RAD, Hercules, CA) before it was concentrated.

### *In vitro* motility assays

G-actin was polymerized in F-buffer (0.1 M KCl, 2 mM MgCl_2_, 1 mM DTT, 1 mM ATP and 10 mM Hepes, pH 7.4) for 1 h at 22°C, and was diluted to 1 μM with NF buffer (25 mM KCl, 2 mM MgCl_2_, 10 mM DTT and 20 mM Hepes, pH 7.4). Diluted actin filaments were incubated with 1 μM rhodamine phalloidin (Invitrogen, Waltham, MA) or 1 μM phalloidin (Wako, Osaka, Japan) for 1 h at 22°C.

*In vitro* actomyosin motility assays, in which actin filaments move on the HMM of skeletal muscle myosin II or full length *Dictyostelium* myosin II, were performed according to the method of Kron and Spudich (Kron and Spudich, 1986), using nitrocellulose-coated flow chambers. In the case of *Dictyostelium* myosin II, full length myosin in 10 mM Hepes, pH 7.4, 200 mM NaCl, 1 mM EDTA, 10 mM DTT was allowed to adhere to the surface, and then incubated with 0.5 mg/ml T166E recombinant myosin light chain kinase (Smith et al., 1996) for 5 min in NF buffer containing 0.5 mM ATP and 10 mg/ml BSA at room temperature. In the case of muscle HMM and myosin filaments, HMM in NF buffer or intact myosin II in 10 mM Hepes, pH 7.4, 50 mM NaCl, 4 mM MgCl_2_, and 10 mM DTT was allowed to adhere to the nitrocellulose surface, followed by blocking with NF buffer containing 10 mg/mL BSA. Rhodamine phalloidin-stabilized actin filaments were then bound to myosin II on a nitrocellulose-coated glass surface in a flow chamber filled with NF buffer containing 10 mg/ml BSA. Movement of actin filaments was initiated by injecting twice the chamber volume of MA buffer (25 mM KCl, 2 mM MgCl_2_, 10 mM DTT, 3 mg/ml glucose, 1.2 μM glucose oxidase, 0.15 μM catalase, 10 mg/ml BSA and 20 mM Hepes, pH 7.4) containing various concentrations of ATP, ADP and Rng2CHD, into the flow chamber. Fluorescence of rhodamine phalloidin was imaged with an EMCCD camera (iXon X3, Andor, Belfast, UK) on a fluorescent microscope (IX-71; Olympus, Tokyo, Japan) equipped with a Plan Apo 100X, 0.9 NA objective lens (Nikon) at a frame rate of 4 fps. The images were processed with Image J (Schneider et al., 2012). For each condition, more than 100 filaments longer than 1.5 µm were randomly selected, and their movements were tracked by MTrackJ, a plugin for Image J (Meijering et al., 2012).

The actomyosin II motility assay in the presence of GFP-Rng2CHD was performed basically as indicated above, with several modifications. Rhodamine phalloidin-stabilized and unlabeled phalloidin-stabilized actin filaments were mixed at a 1:1 concentration in order to observe GFP fluorescence without the interference of rhodamine fluorescence. Fluorescence micrographs were taken with an EMCCD camera (iXon X3) on a TIRF microscope (IX-71) equipped with a UApo N 100X, 1.49 NA objective lens (Olympus). Laser light of 538 nm for exciting rhodamine and 488 nm for exciting GFP were irradiated alternately at 2 s intervals, and the time-lapse imaging of moving actin filaments and GFP-Rng2CHD dynamics was performed semi-simultaneously. The fluorescence intensity of GFP was quantified by Image J as follows. Five filaments near the center of the images were selected for each condition, and the fluorescence intensity in five frames was measured. The light intensity at five points near the filament was averaged and subtracted from the measured values along the filament as the background for each filament. The values were normalized by the length of each filament.

For *in vitro* motility assays in which actin filaments move on myc-tagged myosin V HMM, HMM molecules were immobilized on a glass surface via anti-c-myc antibody. MA2 buffer (20 mM KCl, 4 mM MgCl_2_, 2 mM ATP, 120 µM calmodulin, 1 mM EGTA, 1 mM DTT, 0.5% methylcellulose, 1 mg/ml BSA and 25 mM imidazole, pH 7.4) was used instead of MA buffer. Mouse calmodulin was expressed in *E. coli* and purified as described previously (Shishido et al., 2009). Fluorescence images were captured with a camera (ORCA-Flash 2.8; Hamamatsu Photonics, Hamamatsu, Japan) on a fluorescent microscope (IX-70; Olympus) equipped with a Plan-Fluor 100X, 1.3 NA objective lens (Nikon, Tokyo, Japan) at a frame rate of 0.5 fps.

### Measurement of dissociation constant

G-actin was polymerized in F-buffer for 1 h at 22°C and incubated with phalloidin at a 1:1 molar ratio for 1 h at 22°C. Phalloidin-stabilized actin filaments that were diluted to 3 μM and various concentrations of Rng2CHD or GFP-Rng2CHD (1, 2, 3, 4, 5, 6 µM) were incubated together in SA buffer (25 mM KCl, 2 mM MgCl_2_, 0.5 mM ATP, 10 mM DTT and 20 mM Hepes, pH 7.4) for 5 min at 22°C, and then centrifuged at 278,800 g for 10 min at 22°C. The supernatants and pellets were subjected to SDS-PAGE. Images of Coomassie brilliant blue (CBB)-stained gels were read into Image J and the concentration of Rng2CHD in each fraction was quantified by densitometry. The dissociation constant (*K_d_*) for Rng2CHD to actin filaments was calculated by fitting plots of [Rng2CHD bound to actin filaments] versus [Rng2CHD free] with the following equation:

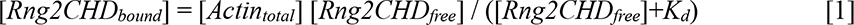

*K_d_* between actin filaments and GFP-Rng2CHD was calculated in the same way.

### Estimation of binding ratio of Rng2CHD and GFP-Rng2CHD to actin filament from *K_d_*

*K_d_* between Rng2CHD and actin filaments is given by:

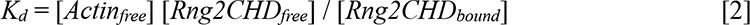

The concentration of actin filaments is extremely low in flow chambers in which Rng2CHD in the buffer interacts with actin filaments immobilized on the substrate and unbound actin filaments were washed away, such as *in vitro* actomyosin motility assays and observations of binding by fluorescence microscopy. Under those conditions, [*Rng2CHD_free_*] can be approximated by the concentration of total Rng2CHD ([*Rng2CHD_total_*]). Therefore, the following approximation holds from Eq. 2:

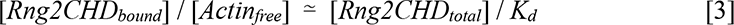

The binding ratios of Rng2CHD and GFP-Rng2CHD to the dilute actin protomers were estimated with this approximation using the value of *K_d_*.

In HS-AFM imaging to measure HHP, unbound actin filaments in solution did not interfere with the imaging, and therefore we were able to include a defined concentration of actin in the observation buffer. In those experiments, the binding ratio was calculated from eq [1].

### High-speed atomic force microscopy

We used a laboratory-built high-speed atomic microscope (HS-AFM) as described previously (Ando et al., 2013). HS-AFM imaging in the amplitude modulation tapping mode was carried out in solution with small cantilevers (BL-AC10DS-A2, Olympus) whose spring constant, resonant frequency in water, and quality factor in water were ∼0.1 N/m, ∼500 kHz, and ∼1.5, respectively. An additional tip was grown, in gas supplied from sublimable ferrocene powder, on the original cantilever tip by electron beam deposition (EBD) using scanning electron microscopy (ZEISS Supra 40 VP/Gemini column, Zeiss, Jena, Germany). Typically, the EBD tip was grown under vacuum (1 - 5 x 10^-6^ Torr), an aperture size of 10 µm, and electron beam voltage of 20 keV for 30 s. The EBD ferrocene tip was further sharpened using a radio frequency plasma etcher (Tergeo Plasma Cleaner, Pie Scientific, Union City, CA) under an argon gas atmosphere (typically at 180 mTorr and 20 W for 30 s). For HS-AFM imaging, the free-oscillation peak-to-peak amplitude of the cantilever (*A_0_*) was set at ∼1.6 - 1.8 nm, and the feedback amplitude set-point was set at ∼0.9 *A_0_*.

Liposomes composed of DPPC/DPTAP (90/10, wt/wt) and mica-supported lipid bilayer were made according to our previous sample preparation protocol (Ngo et al., 2015). We used this positively charged lipid bilayer for gently immobilizing actin filaments in all HS-AFM experiments. In the first set of experiments, we observed the impact of different Rng2CHD binding ratios on the structure of actin filaments at the equilibrium binding states between Rng2CHD and actin filaments. Actin filaments were initially made at a final actin concentration of 20 μM in F-buffer containing 0.1 M KCl, 2 mM MgCl2, 1 mM DTT, 1 mM ATP and 10 mM Hepes-KOH pH 7.4 for ∼1 h at 22°C. For HS-AFM imaging of actin filaments at different Rng2CHD binding ratios, we fixed the final concentration of actin filaments in the AFM observation chamber at 0.59 μM in the observation buffer (25 mM KCl, 2 mM MgCl_2_, 50 µM ATP, 1 mM ADP, 10 mM DTT, and 20 mM Hepes pH 7.4) and calculated the binding ratios at different concentrations of Rng2CHD using a *K_d_* of 0.92 μM (Table Supplement 1). The protein bindings were allowed after an incubation of the mixture in a tube at 22°C for 10 min or longer. The protein mixture (68 μl) was added into the observation chamber, in which the positively charged lipid bilayer was already made, followed by the approaching process of the sample scanner stage. The actin filaments at different Rng2CHD binding ratios were gently immobilized onto this lipid bilayer during sample approaching (∼5-7 min), prior to the HS-AFM imaging. HS-AFM imaging process was performed as described in detail elsewhere (Ando et al., 2013), except an additional use of a recently developed OTI mode (Fukuda et al., 2021). Half helical pitches (HHPs) of actin filaments were analyzed by measuring the distance between the crossover points of two single actin protofilaments along the filaments using a home-built software (UMEX Viewer for Drift Analysis), which allowed us to semi-automatically determine and measure the distance between highest points of two neighboring actin protomers (e.g., HHPs) by making a topographical line profile along actin filaments. Briefly, a cross-sectional profile line was drawn along the long axis of the actin filament with the length of 1 – 3 consecutive half helices. Prior to the analysis, the nonlinearity of the XY piezos was corrected by a nonlinear image scaling, and the image noise was suppressed by a Gaussian smooth filter with the standard deviation of 0.76 nm. The profile was extracted by averaging the signal in a 6 nm band along the filament. To reduce the effect of noise, we set minimum threshold pitch values of 3 and 20 nm for the actin protomer and HHP, respectively. The measured HHP data were copied into an Excel datasheets for statistical analysis.

In the second set of experiments, we analyzed the impact of Rng2CHD on the binding of S1 to actin filaments. G-actin was polymerized in F-buffer for 1 h at 22°C, and then the buffer on the sample stage was replaced by 2 μl of 20 μM actin filaments in HSAFM buffer (25 mM KCl, 2 mM MgCl_2_, 50 μM ATP, 1 mM ADP, 10 mM DTT, 20 mM Hepes, pH 7.4). After incubation for 10 min at 22°C, the surface of the sample stage was rinsed with 20 μl of HSAFM buffer to remove free actin filaments. Subsequently, the surface of the sample stage was immersed in 60 μl of HSAFM buffer in the observation chamber of the HS-AFM. Observations of the transient binding of muscle S1 to actin filaments in the presence of ATP and ADP were performed under the following two conditions: (1) S1 diluted in HSAFM buffer was added to the observation chamber to a final concentration of 500 nM; (2) 12 nM Rng2CHD was allowed to interact with actin filaments in the observation chamber for 15 min at 22°C, and then 20 µM S1 in HSAFM buffer was added to the observation chamber to a final concentration of 500 nM. AFM images were obtained at a scan speed of 0.5 s per field of view, and were then visualized by Kodec4, our laboratory-built software (Ngo et al., 2015). Images scanned between 1 min and 2 min after the addition of S1 or ATP were analyzed, and the events of transient binding of S1 molecules to actin filaments were visually counted.

### Electron microscopy

G-actin was polymerized in F-buffer for 1 h at room temperature, and 1 μM actin filaments were mixed with various concentrations of Rng2CHD in EM buffer (25 mM KCl, 4 mM MgCl_2_, 1 mM DTT, 0.1 mM ATP and 10 mM imidazole, pH 7.4) at room temperature. A small volume of each sample was placed on a carbon-coated grid for 30 s (40 nM Rng2CHD), 2 min (200 nM Rng2CHD) or 4 min (1 μM Rng2CHD) after mixing. The samples were negatively stained with 1% uranyl acetate, and observed in a transmission electron microscope (Tecnai F20; FEI, Hillsboro, OR). Electron micrographs were recorded with a Gatan ORIUS 831 CCD camera (Pleasanton, CA), adjusted for contrast and Gaussian-filtered using Adobe Photoshop.

### Myosin II S1 ATPase measurements

Actin-activated S1 ATPase was measured using malachite green (Kodama et al., 1986). G-actin was polymerized in F-buffer for 1 h at 22°C. The solution was centrifuged at 278,800 g for 10 min at 22°C and actin filaments in the pellet were resuspended in NF buffer. This procedure was repeated once more to minimize the amount of carried-over phosphate. Actin filaments and various concentrations of Rng2CHD (0, 0.33, 0.82, 1.9, 5.0 μM) were mixed in NF buffer containing 2 mM ATP, then incubated for 10 min at 25°C. The reaction was started by the addition of S1, and phosphate released at 0, 2, 4, 6 and 8 min was measured. The final concentrations of actin and S1 were 24 µM and 50 nM, respectively.

### Co-sedimentation assay of actin filaments and S1 with Rng2CHD

For the co-sedimentation assays in the presence of ATP, G-actin was polymerized in F-buffer for 1 h at 22°C, then incubated with phalloidin at a 1:1 molar ratio for 1 h at 22°C. For the co-sedimentation assays in the presence of ADP, the solution of actin filaments was centrifuged at 278,800 g for 10 min at 22°C, and the pelleted actin filaments were resuspended in NF buffer. This procedure was repeated once again to minimize the amount of carried-over ATP before the addition of phalloidin. The following two samples were prepared for both experiments: (1) 3 μM actin filaments were incubated with 2 μM S1 for 5 min at 22°C; (2) 3 μM actin filaments were incubated with 2 µM Rng2CHD for 10 min at 22°C, and after adding S1, were incubated for 5 min at 22°C. Each sample was prepared in S-ATP buffer (25 mM KCl, 2 mM MgCl_2_, 2 mM ATP, 10 mM DTT, 10 mM phosphoenolpyruvate, 10 units/ml pyruvate kinase and 20 mM Hepes, pH 7.4) or S-ADP buffer (25 mM KCl, 2 mM MgCl_2_, 2 mM ADP, 10 mM DTT and 20 mM Hepes, pH 7.4). After incubation, each sample was centrifuged at 278,800 g for 10 min at 22°C. The supernatants and pellets were subjected to SDS-PAGE. Images of CBB-stained gels were read by and into Image J and the concentration of S1 in each fraction was quantified by densitometry.

### Fluorescence microscope-based binding assay

Binding of HMM-GFP to actin filaments was observed as follows. G-actin was polymerized in FF-buffer (50 mM KCl, 2 mM MgCl_2_, 0.5 mM EGTA, 1 mM DTT, 20 mM PIPES, pH 6.5) containing 0.2 mM ATP for 2 h at 22°C. Actin filaments and Alexa 647 phalloidin (Invitrogen) were mixed at a molar ratio of 20:1 and incubated overnight on ice. The surface of each coverslip was covered with a positively charged lipid bilayer and was used to construct flow chambers as described previously (Ngo et al., 2015), except that the weight ratio of 1,2-dipalmitoyl-sn-glycero-3-phosphocholine (DPPC; Avanti Polar Lipids, Alabaster, AL) and 1,2-dipalmitoyl-3-trimethylammonium-propane (DPTAP; Avanti Polar Lipids) was 17:3 (Hirakawa et al., 2017; Hosokawa et al., 2021). Alexa 647 phalloidin-stabilized actin filaments diluted in FF-ATP buffer (FF buffer containing 0.5 µM ATP) were introduced into the flow chamber to loosely bind to the positively charged lipid layer. HMM-GFP and Rng2CHD diluted in FF-ATP buffer were then introduced to the flow chamber. Alternatively, FF-ATP buffer in the above procedures was replaced with FF buffer for the assays in the nucleotide-free state. Fluorescence of Alexa 647 and GFP was imaged with a fluorescence microscope (ECLIPSE E600, Nikon) equipped with an ARUGUS-HiSCA system (Hamamatsu Photonics). Images were captured using a 100x objective lens (CFI Plan Apo Lambda 100x Oil, NA 1.45; Nikon).

## Acknowledgments

We thank Dr. Atsuko H. Iwane and Dr. To shio Ya nagida for the gift of the virus to express HMM of myosin V. This work was supported in part by Bio-SPMs Collaborative Research of WPI Nano Life Science Institute, Kanazawa University, and Grants-in-aid from the Ministry of Education, Culture, Sports, Science and Technology to KT (No. 24370069 and 24117008), KN (No. 22019004), MT (No. 24770177) and TU (No. 24370069 and 24117008).

## Author contributions

NK, KT, ON, TF, TA, KN and TU conceived and supervised the study, YH, MT, KXN, KT, KN and TU designed the experiments, YH, MT, KXN, TI, MY, ABB,KH, AY and TU performed the experiments, YH, MT, KXN, MY, ABB, KU, KH and TU analyzed the data, YH and TU wrote the manuscript draft and KXN, KH, NK, KT, TF, MT and KN revised the manuscript.

## Conflicts of interest

The authors declare no conflicts of interest.

**Videos** (the video files can be downloaded at https://www.dropbox.com/sh/r1qvgkjlxjzzvxk/AAACHWH0ZeRzxgnW1eemp2qGa?dl=0)

**Video 1.** Movement of actin filaments on surfaces coated with full length *Dictyostelium* myosin II in the absence (left) or the presence of 200 nM (middle) and 1 µM (right) Rng2CHD. The concentration of ATP was 1 mM. Playing speed is 1x.

**Video 2.** Movement of actin filaments on surfaces coated with HMM of rabbit skeletal muscle myosin II in the absence (left) or the presence of 50 nM (middle) and 200 nM (right) Rng2CHD. The concentration of ATP was 1 mM. Playing speed is 1x.

**Video 3.** Movement of actin filaments on surfaces coated with filaments of rabbit skeletal muscle myosin II in the absence (left) or presence of 100 nM Rng2CHD (right). The concentration of ATP was 0.5 mM. Playing speed is 1x.

**Video 4.** Movement of actin filaments on surfaces coated with HMM of rabbit skeletal muscle myosin II in the presence of 200 nM Rng2CHD and in the presence of 1 mM ATP (left) or 0.2 mM ATP and 1 mM ADP (right). Playing speed is 7.5x.

**Video 5.** Movement of actin filaments on surfaces coated with HMM of mouse myosin V in the absence (left) or the presence of 330 nM Rng2CHD (rigbt) Rng2CHD. The concentration of ATP was 2 mM. Playing speed is 1x.

**Video 6.** Transient binding of Rng2CHD. Real time HS-AFM observation of 0.59 µM actin filaments interacting with 0, 0.25, 0.85 and 5.7 µM Rng2CHD. The estimated binding ratios under those conditions are 0, 15%, 40% and 85%, respectively. Red and green arrowheads denote sparse binding of Rng2CHD molecules and some typical Rng2CHD clusters, respectively. Imaging rate was 2 frame/s and the playing speed is 15x. Bars: 25 nm. For details, see Figure 4.

**Video 7.** Distortion of helical structures of actin filaments and separation of protofilaments induced by high concentration of Rng2CHD. Imaging rate was 1 frame/s, and the playing speed is 10x. Bars: 25 nm. For details, see Figure Supplement 3.

## Supplementary Materials

### Supplementary information 1. Estimation of binding ratio of GFP-Rng2CHD along actin filaments based on *Kd*

Densitometric scanning of a cosedimentation gel shown in Figure 3D resulted in the binding curve shown below. Fitting of the data points, shown in dotted line, yielded the *K_d_* value of 3.4 µM.

The table below shows the binding densities of Rng2CHD needed to achieve several degrees of motility inhibition on muscle HMM, calculated based on *K_d_* (for Rng2CHD and GFP-Rng2CHD) and also directly from the fluorescence intensities of GFP-Rng2CHD.

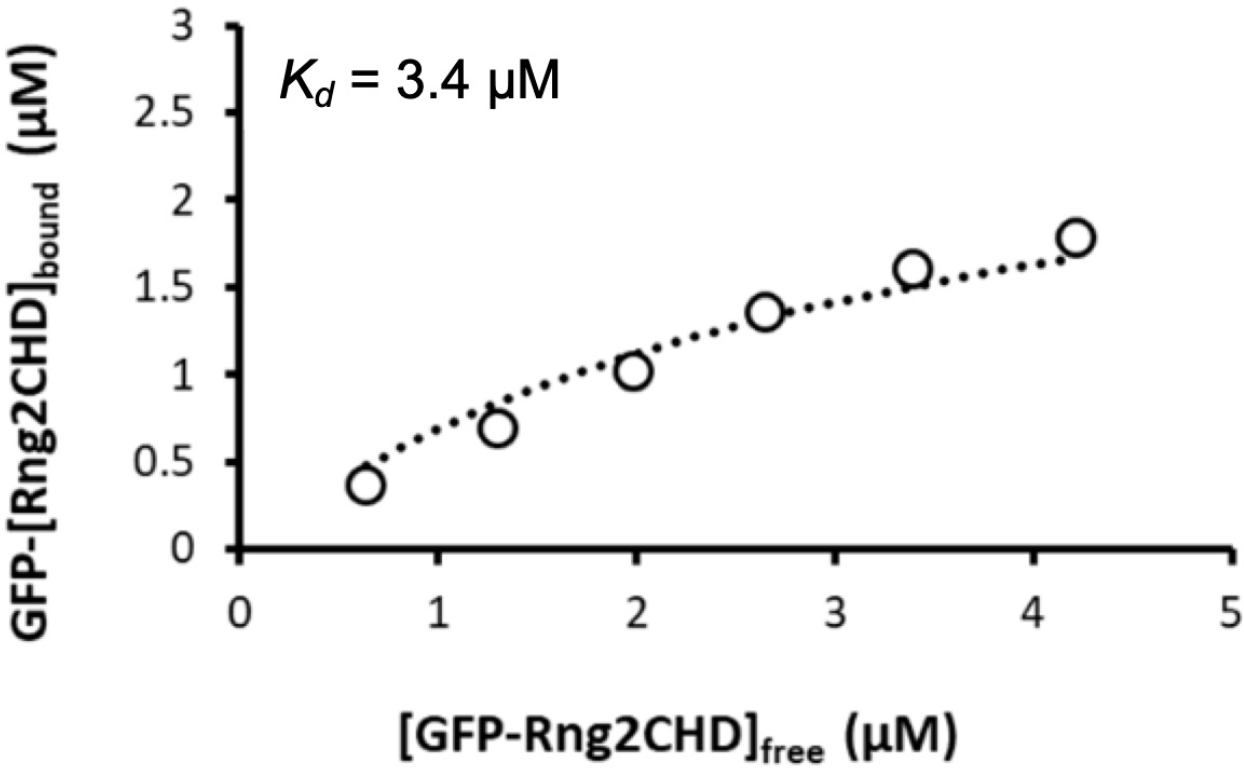

**Table.**
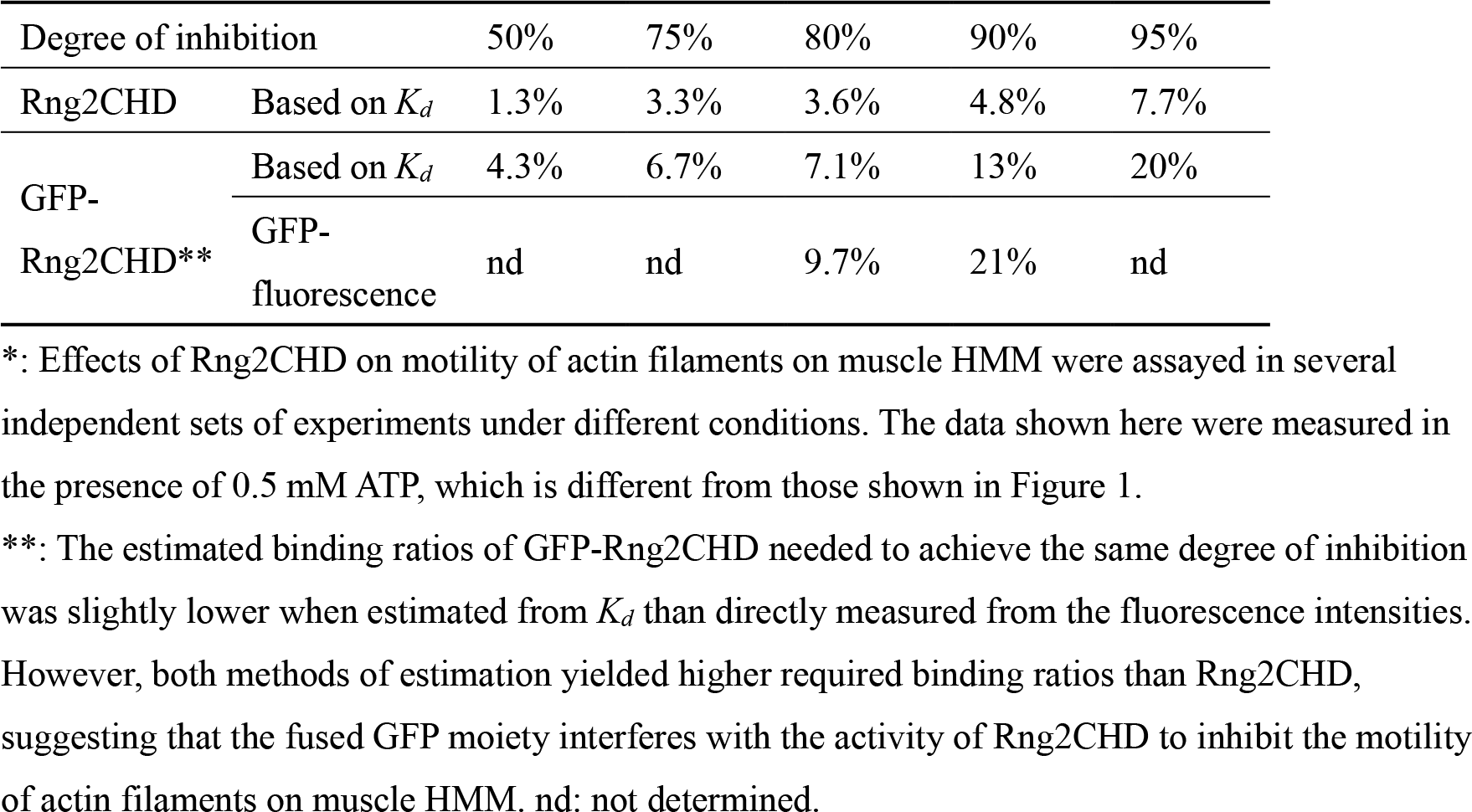
Binding ratio of Rng2CHD needed to achieve various degrees of motility inhibition*.

### Supplementary information 2. Possible mechanisms for differential responses of skeletal muscle myosin II and *Dictyostelium* non-muscle myosin II to Rng2CHD

Although actin movements by both skeletal muscle and *Dictyostelium* myosin IIs were inhibited by Rng2CHD, the apparent modes of inhibition were very different. On muscle myosin II-coated surfaces, actin movements were inhibited by relatively low concentrations of Rng2CHD and were virtually immobilized at 200 nM Rng2CHD (Figure 1A, Video 2 and Figure Supplement 1). In contrast, actin filaments moving on *Dictyostelium* myosin II-coated surfaces appeared to lose affinity with the surface in the presence of medium concentrations of Rng2CHD without slowing significantly, and they diffused away from the surface in the presence of high concentrations of Rng2CHD (Figure 1B and 1C and Video 1). This apparent difference can be explained, at least in part, by two quantitative differences between muscle and *Dictyostelium* myosin IIs within the framework that Rng2CHD impairs the generation of active force by affecting the transition from the A•M•ADP•Pi complex to the A•M•ADP complex and/or the affinity of the A•M•ADP complex.

The first quantitative difference is the affinity for actin in the so-called weakly bound state. *Dictyostelium* myosin II has only three positive charges in loop 2, while muscle myosin II has five. Loop 2 is a major electrostatic actin binding site in the weakly bound state, and the number of positive charges in loop 2 determines the affinity for actin in the weakly bound state (Furch et al., 1996). The resistive load by increasing the number of weakly-bound muscle myosin II motors has been shown to slow and ultimately stop the movement propelled by active muscle HMM (Warshaw et al., 1990). In contrast, the weaker affinity of *Dictyostelium* myosin II in the weakly bound state would lead to smaller resistive load, which could result in an apparently very different outcome when the generation of active force is reduced by Rng2CHD. Moreover, the weaker weakly bound state with *Dictyostelium* myosin II would allow the actin filaments to detach from the myosin-coated surface when tethering by strongly-bound force-generating interactions is shortened or weakened by Rng2CHD.

The second quantitative difference that could contribute to the apparent difference in sensitivities of the two myosin IIs to Rng2CHD is the longer duration of the strongly-bound, force-generating A•M•ADP complex of *Dictyostelium* myosin II, and this mechanism is supported by the fact that the movement by muscle HMM became less sensitive to Rng2CHD in the presence of 0.2 mM ATP and 1 mM ADP (Figure 1B, Video 3 and Figure Supplement 1). We presume that in the presence of intermediate concentrations of Rng2CHD (i.e., 100-200 nM), many, but not all, of the productive force-generating events are inhibited, so that the remainder of the productive interactions are insufficient to move actin filaments against the resistive load imposed by weakly-bound crossbridges of muscle myosin II. However, the small number of force-generating crossbridges may be sufficient to move actin slowly when duration of the tension-bearing A•M•ADP complexes is extended. In the case of *Dictyostelium* myosin II, the resistive load by weakly-bound crossbridges is smaller, and the duration of the tension-bearing A•M•ADP complexes is longer, so that Rng2CHD-affected actin filaments move on *Dictyostelium* myosin II without slowing significantly, and eventually detach from the myosin-coated surfaces.

Rng2CHD-induced weaker affinity of the A•M•ADP complex could have an interesting consequence in the case of *Dictyostelium* myosin II. Shortening of the lifetime of the affected A•M•ADP complexes may accelerate the movement within a certain range of Rng2CHD concentrations when the ensemble of the active force is sufficient to overcome the resistive load. This may be the reason why the average actin velocity was slightly faster in the presence of 50 nM Rng2CHD than in its absence (Figure 1B).

Although the combination of the above two quantitative differences can explain, at least in part, the observed apparent difference in the response of the two myosin IIs to Rng2CHD, we cannot rule out the possibility that the Rng2CHD affects the motilities of muscle and *Dictyostelium* myosin IIs in a manner not easily predicted within the framework of the standard swinging lever-arm model.

### Supplementary information 3. Comparison of the actin sliding speed by myosin II *in vitro*, and the contraction speed of contractile rings *in vivo*

The contraction speed of contractile rings (CRs) is much slower than the sliding velocity of actomyosin II *in vitro*. For instance, CRs in *S. pombe* contract at a rate of 0.5 μm/min circumferentially (Pelham and Chang, 2002), while actin filaments on *S. pombe* myosin II-coated surfaces move at a rate of 30 μm/min *in vitro* (Lord and Pollard, 2004). In *Dictyostelium*, the circumferential contraction velocity of the CR is 0.18 μm/s (Zang et al., 1997), whereas the sliding velocity of actin filaments on *Dictyostelium* myosin II is 1.1 μm/s (Figure 1B) or 1.4 µm/s (Kron and Spudich, 1986) *in vitro*, more than 5-fold faster than the circumferential speed of CR contraction.

A direct comparison of *in vitro* sliding speed and contraction speed is difficult for two reasons. The first aspect to complicate the quantitative comparison of the two speeds is the geometry of CRs. In the case of the simplest model CR that consists of a single contractile unit, shown below, the circumference of the CR would contract at 2x the speed of unitary actomyosin sliding speed. If CR consists of a series of multiple contractile units, as would be expected for real CRs, the circumferential speed of contraction would be further multiplied by the number of contractile units. Thus, the 60-fold (*S. pombe*) or 5-fold (*D. discoideum*) difference between the *in vivo* and *in vitro* speeds is vastly an underestimate of the real difference.

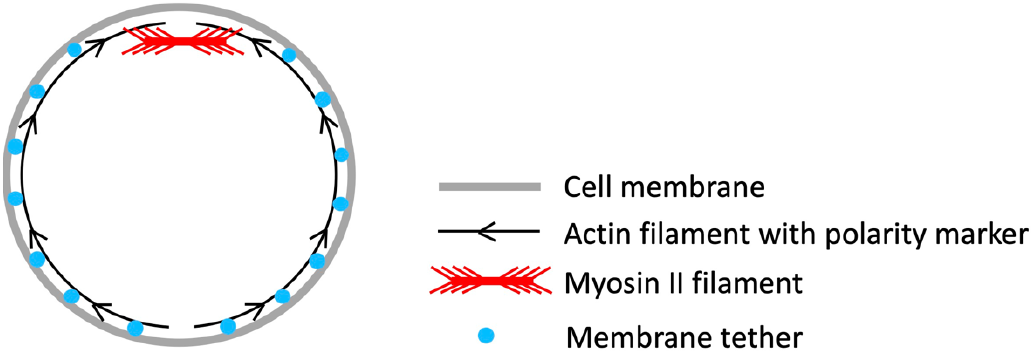

The second issue to be considered is the possible slowing of the actomyosin movement due to large load to contract CRs. Quantitative assessment of this possibility is difficult, but mutant *Dictyostelium* cells lacking myosin II on substrates can divide at a rate only 2-fold slower than the wild type cells (Zang et al., 1997), implying that at least in the case of adherent *Dictyostelium* cells, a very large force is not necessary to drive contraction of the CRs.

Based on these reasons, we speculate that contraction of the CR is regulated in an inhibitory manner in cells with and without a cell wall.

---

**Figure Supplement 1.**
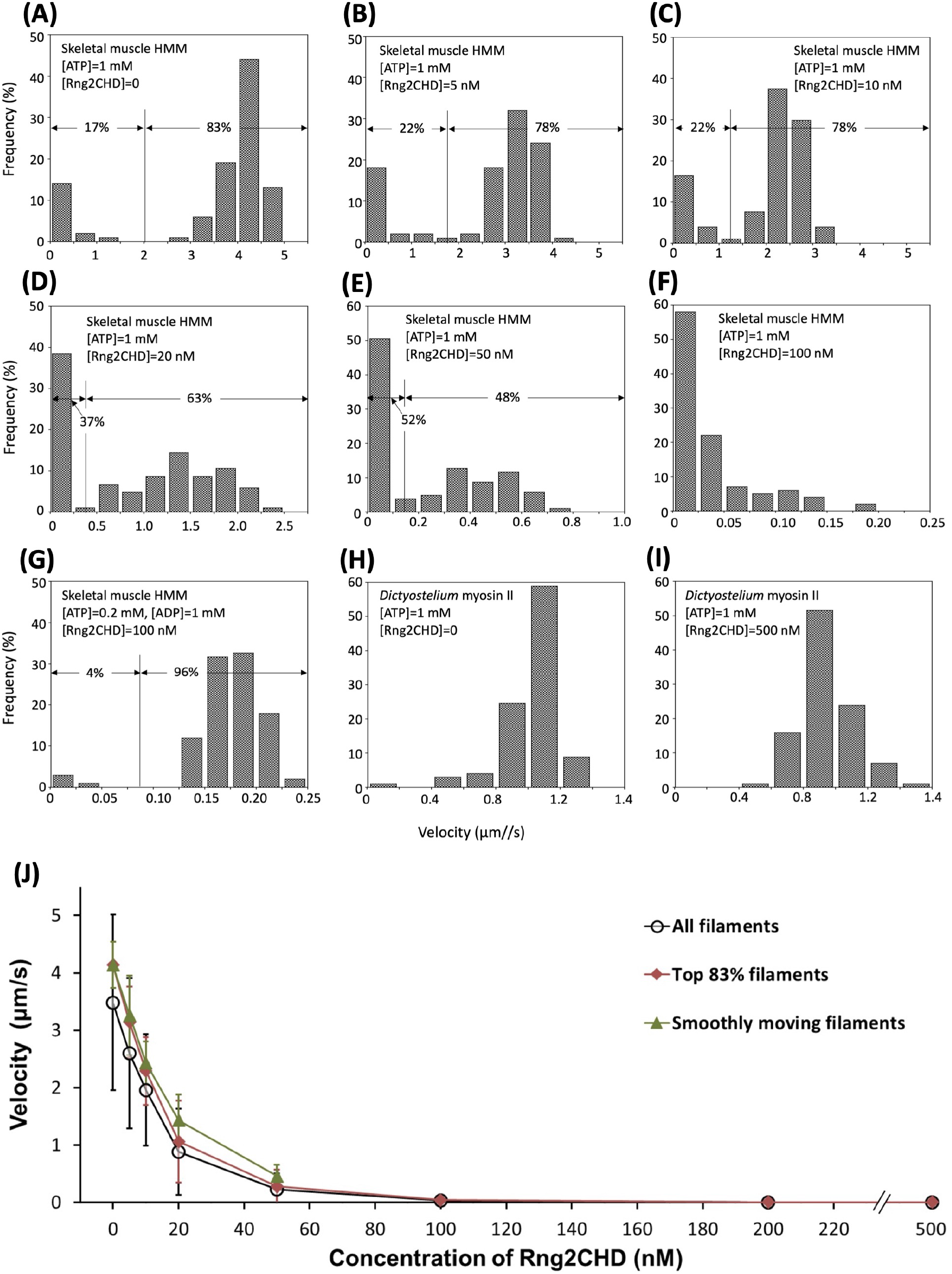
Detailed analyses on inhibition of actin-myosin II movements by Rng2CHD. Contaminating “bad” myosin heads that do not release actin filaments in the presence of ATP would act as break and slow or stall the actin movements in *in vitro* motility assays. In the experiment shown in Figure 1A, a non-negligible fraction of the filaments were immobile in the absence of Rng2CHD. For the evaluation of motor activity of myosin *in vitro*, researchers often ignore those immobile filaments, visually select smoothly moving filaments and measure their velocities. This was not appropriate in this experiment, since Rng2CHD decreased the mean velocity by increasing the fraction of immobile filaments, in addition to slowing the smoothly moving filaments. This is the reason why we presented mean velocity of all actin filaments, including the immobile ones, under each experimental condition in Figure 1. In this Figure Supplement, we separately show contributions of the two effects, i.e., increase of the stalled filaments and slowing of the smoothly moving filaments, by Rng2CHD. In (**A-E**), distribution of velocities of all measured filaments are shown. In the absence of Rng2CHD, velocities on muscle HMM in the presence of 1 mM ATP had two distinct populations, one near zero, and the other centered around 4.1 µm/s. The fraction of the “immobile” filaments was 17%, and the faster one, which corresponds to smoothly moving filaments as judged by visual inspection of the traces, was 83%. The mean velocity of the immobile fraction was 0.22 ± 0.32 µm/s (mean ± SD, N=18), while that of the faster “smoothly moving” filaments was 4.1 ± 0.4 µm/s (N=83), with the mean velocity of all filaments of 3.5 ± 1.5 µm/s. Two distinct velocity distributions were recognizable up to the Rng2CHD concentration of 50 nM, when the fraction of the immobile filaments increased to 52% and the mean velocity of the smoothly moving filaments decreased to 0.46 ± 0.2 µm/s, with the mean velocity of all filaments of 0.23 ± 0.28 µm/s (**E**). At the Rng2CHD concentration of 100 nM or higher, the velocities of moving filaments became so slow that the distinction of two velocity fractions became impossible (**F**). Intriguingly, in the presence of 0.2 mM ATP and 1 mM ADP, most of the filaments were moving smoothly, albeit slowly, at 0.17 ± 0.04 µm/s even in the presence of 100 nM or 200 nM (not shown) Rng2CHD, with virtually no immobile filaments (4%) (**G**). Similarly, most of the filaments were moving smoothly on *Dictyostelium* myosin II, both in the absence (**H**) and presence of Rng2CHD (**I**). **J** shows mean ± SD actin velocities on muscle HMM in the presence of 1 mM ATP that were calculated by three different methods. Velocities of smoothly moving filaments were calculated up to the Rng2CHD concentration of 50 nM. The SDs are fairly large for all actin filaments (43% of the mean in the absence of Rng2CHD), but are reasonably small for smoothly moving filaments (10% of the mean). In the absence of Rng2CHD, 17% of the filaments were immobile, implying that immobile fraction in excess of 17% in the presence of Rng2CHD is due to the inhibitory effect of Rng2CHD. With this premise, we also calculated the mean velocities of the faster 83% filaments in each condition. This set of data may more accurately represent the overall inhibitory effect of Rng2CHD on motility by muscle HMM. <u>Related to</u> Figure 1.

**Figure Supplement 2.**
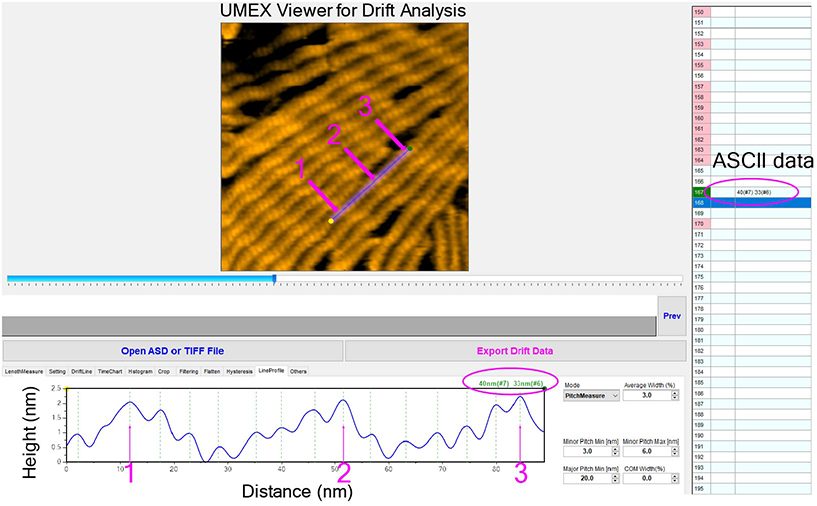
A screenshot demonstration of the peak determination and measurement of half helical pitch (HHP) of actin filaments using a semi-automated software UMEX Viewer for Drift Analysis. The typical threshold parameters were set to unambiguously determine the positions of actin protomers and measure HHPs along actin filaments, as described in detail in Materials and Methods. Magenta arrows denote the highest points (e.g., crossover points) automatically determined on a line drawn along an actin filament in the middle of the field (a major pitch min was set at 20 nm). Green dashed lines denote the positions of the highest points of individual actin protomers (the minor pitch min and max were set at 3 and 6 nm, respectively). For example, the two HHPs were measured between three adjacent purple arrows (peak 1, 2, 3) along an actin segment. The HHP and number of actin protomers counted per one HHP are shown simultaneously, as referred to the data marked by magenta oval. Typically, 20 – 30 actin filaments were selected and HHPs were measured for ∼20 consecutive time frames. To minimize the measurement error of HHP, the straight actin segments were selected and the line profile (average width of 3%) was normally drawn to measure 1-3 consecutive half helices. <u>Related to</u> Figure 4.

**Figure Supplement 3.**
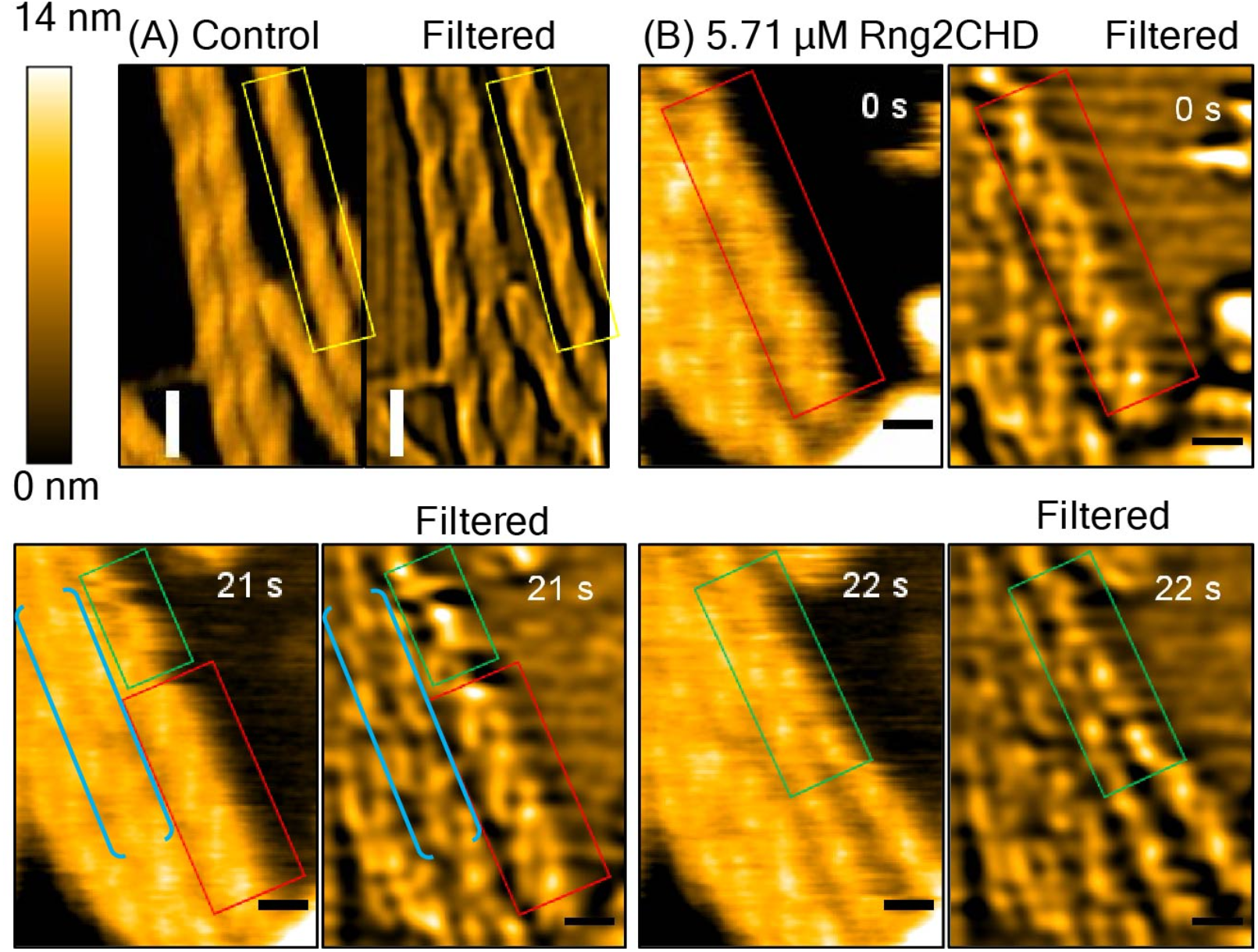
Distortion of helical structures of actin filaments and separation of protofilaments induced by high concentration of Rng2CHD. Control actin filaments (**A**) and actin filaments incubated with 5.7 µM Rng2CHD (**B**) were imaged by HS-AFM at different time points. Yellow rectangles denote control actin segments with compact and normal helical structures. Red rectangles denote actin filaments bound with Rng2CHD with abnormal helical structures. Green rectangles denote two single actin protofilaments separated from each other. Brackets denote actin segments that contain both abnormal helical structures and two separated protofilaments. Filtered AFM images were made to visualize more clearly helical structures of actin filaments by applying simultaneously Laplacian and Gaussian filters (sigma of 20). Bars: 25 nm. A stack of these images is shown in Video7. <u>Related to</u> Figure 4.

**Figure Supplement 4.**
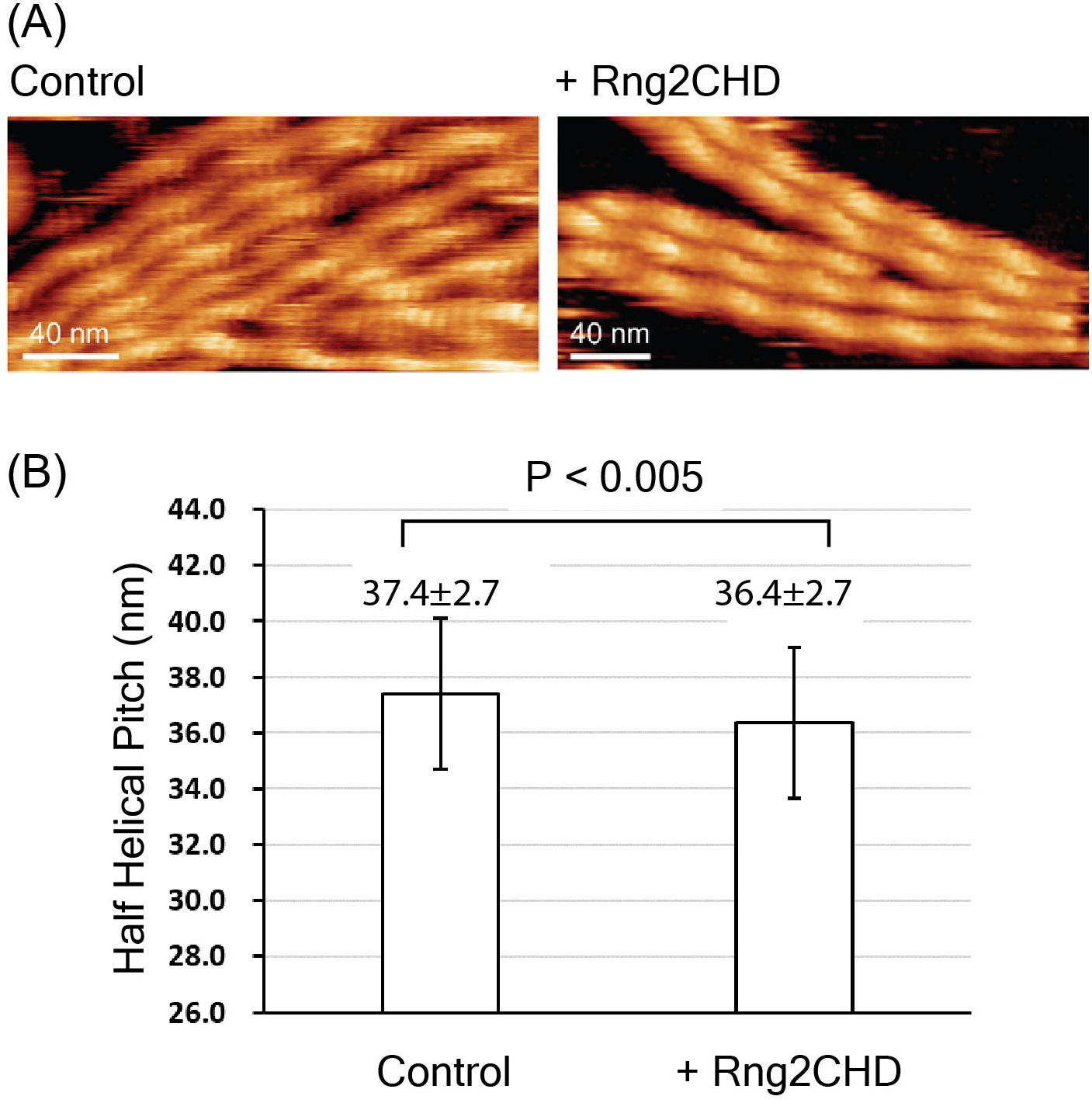
High-resolution AFM topography of actin subunits in control and Rng2CHD-treated actin filaments acquired by FM-AFM. (**A**) Typical high-resolution AFM topography image of actin subunits in control and Rng2CHD-treated filaments used for analyzing HHP. (**B**) Histogram of the HHP of control and Rng2CHD-treated actin filaments acquired by FM-AFM. <u>Related to</u> Figure 4. **Methods**. We used a laboratory-built frequency modulation atomic force microscopy (FM-AFM) system operating in liquid environments equipped with an ultra-low noise cantilever deflection sensor (Fukuma et al., 2005; 2006). The photothermal excitation technique was used to drive the cantilever oscillation (Fukuma et al., 2012). AFM scanning was controlled by a commercial SPM controller (ARC2, Asylum Research, Santa Barbara, CA), and the oscillation amplitude of the cantilever was kept constant using a controller (OC4, SPECS, Berlin, Germany). AFM was operated in constant frequency shift Δ*f* mode, where the tip-sample distance was adjusted such that Δ*f* was kept constant. AFM data were acquired using an AFM cantilever (240AC-NG, OPUS, Sofia, Bulgaria) with nominal spring constants of 2 N/m and a nominal tip radius of ≤7 nm. After each measurement, the actual spring constant of each cantilever was determined using the thermal noise method (Hutter et al., 1993). The obtained AFM images were processed by using WSXM or Gwyddion software. G-actin was polymerized in FM-AFM buffer (40 mM KCl, 1 mM MgCl_2_, 1 mM DTT, 0.5 mM EGTA, 1 mM ATP and 20 mM PIPES, pH 6.8) for 1 h at 22°C, and then 2.1 μM actin filaments were gently mixed with 0.32 µM Rng2CHD and incubated at room temperature for 5 to 10 min. Immediately, the control and Rng2CHD-treated actin filaments were deposited on a positively charged lipid bilayer (DPPC/DPTAP, 25/75 wt/wt) for 15 min, subsequently washed with 0.25x FM-AFM buffer 2 or 3 times prior to imaging in 0.25x FM-AFM buffer.

**Figure Supplement 5.**
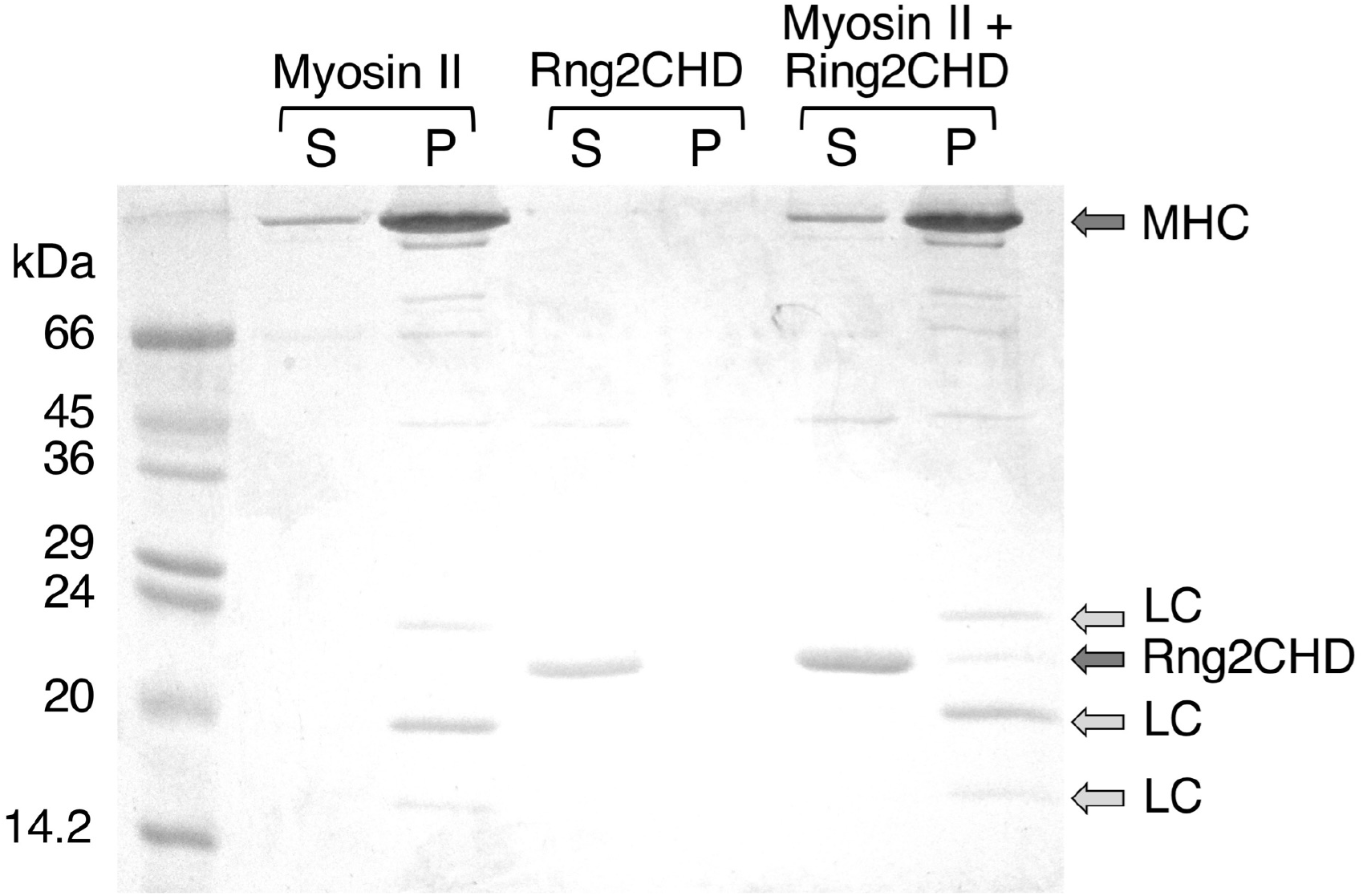
Co-sedimentation of Rng2CHD with skeletal muscle myosin II filaments. Skeletal muscle myosin II filaments, Rng2CHD and a mixture of the two proteins were centrifuged at 30,000 rpm for 10 min at 22°C in a Beckman TLA100.1 rotor, and the supernatant and pellet fractions were analyzed by SDS-PAGE. The concentration of myosin II and Rng2CHD was 2 µM, and the buffer was 10 mM Hepes pH 7.4, 25 mM KCl, 4 mM MgCl_2_, 0.2 mM ATP and 1 mM DTT.

**Figure Supplement 6.**
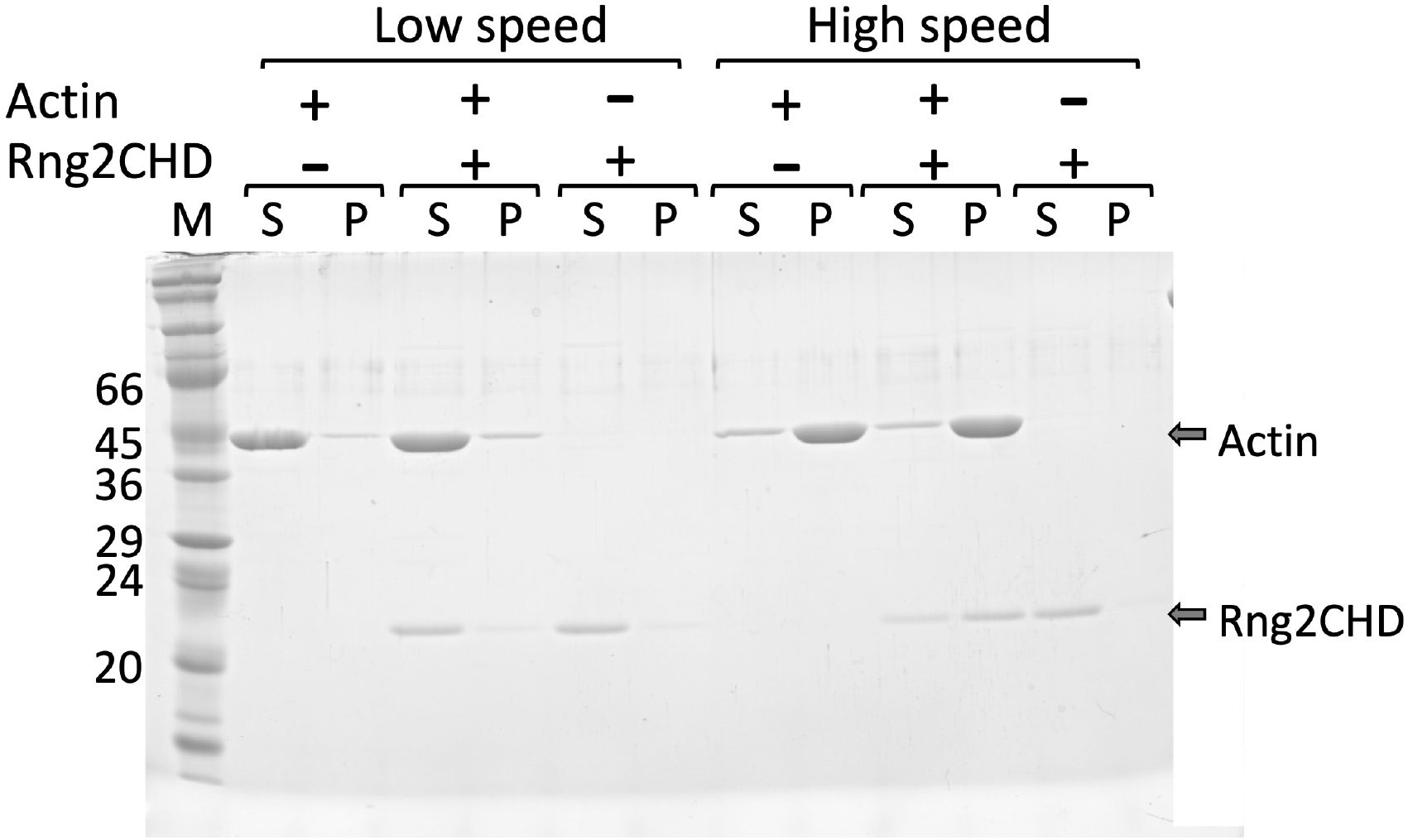
Co-sedimentation of Rng2CHD with actin filaments by low and high speed centrifugation. Mixtures of 3 µM actin filaments and 1.5 µM Rng2CHD in KMEI (0.1 M KCl, 2 mM MgCl_2_, 0.5 mM EGTA, 0.5 mM DTT, 0.2 mM ATP, and 10 mM imidazole, pH 7.5) were incubated for 1 h at 25°C. They were then centrifuged at 20,000 x *g* (low speed) for 15 min or at 200,000 x *g* (high speed) for 20 min, and pellet and supernatant fractions were analyzed by SDS-PAGE. Rng2CHD cosedimented with actin filaments after ultracentrifugation, but neither protein was pelleted by low speed centrifugation. Under the same condition, His-Rng2CHD–bundled actin filaments and both proteins were pelleted by low speed centrifugation (Takaine et al., 2009).

**Table Supplement 1.**
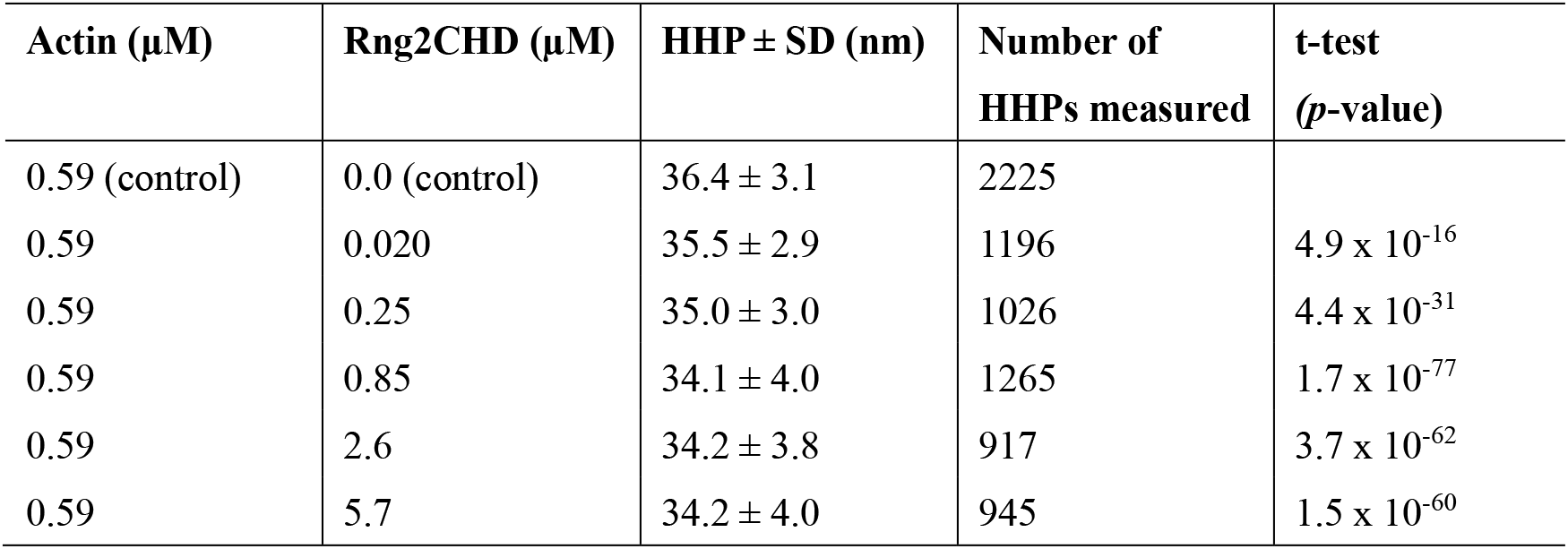
Half helical pitch (HHP) of actin filaments incubated with different Rng2CHD concentrations at the equilibrium state (*K_d_* of Rng2CHD = 0.92 µM). The value is a mean HHP ± SD. The differences between the mean HHP of control actin filaments (0 µM Rng2CHD) and those incubated with different Rng2CHD concentrations were statistically significant (*, p ≦ 0.001, two independent populations *t*-test). **<u>Related to</u>** Figure 4.

## Notes

### Competing Interest Statement

The authors have declared no competing interest.

### Summary of Updates

\# Effects of Rng2CHD on motility by non-muscle myosin II (Dictyostelium myosin II) was investigated. Response of Dictyostelium myosin II to Rng2CHD was very different from that of muscle myosin II, but the difference can be explained on the basis of quantitative differences between these two myosin IIs. # Structural changes of actin filaments induced by Rng2CHD were investigated using HS-AFM. The results demonstrated that Rng2CHD induces cooperative conformational changes to actin filaments that are accompanied by supertwisting of the actin helix, at substiochiometric binding densities.

